# Threshold-Free Network-Oriented Statistics in Neuroscience

**DOI:** 10.1101/2024.01.03.574016

**Authors:** Zexuan Hao, Pei Wang, Xiaoyu Xia, Yu Pan, Weibei Dou

**Affiliations:** Department of Electronic Engineering, Beijing National Research Center for Information Science and Technology (BNRist), Tsinghua University, Beijing 100084, China; Department of Statistics, Miami University, Oxford, OH 45056, USA; Senior Department of Neurosurgery, The First Medical Center of PLA General Hospital, Beijing 100853, China; Department of Neurosurgery, The Seventh Medical Center of Chinese PLA General Hospital, Beijing 100700, China; Department of Rehabilitation Medicine, School of Clinical Medicine, Beijing Tsinghua Changgung Hospital, Tsinghua University, Beijing 102218, China

**Keywords:** Multiple testing, Threshold-free statistics, Network, Graph, Connectivity, Statistical power

## Abstract

Network neuroscience has emerged as an indispensable tool for studying brain structure and function. Currently, the network-based statistic (NBS) procedure is widely used for dealing with massive multiple testing/comparison problems in brain networks. However, the NBS requires choosing a hard cluster-forming threshold, lacking objective rules. A powerful and flexible statistical framework is urgently needed with growing interest in finer-grained network explorations across modalities and scales. Here, we introduce a permutation-based framework—”Threshold-Free Network-Oriented Statistics” (TFNOS). It integrates two “threshold-free” pathways: traversing all cluster-forming thresholds (TT) and using predefined clusters (PC). The TT procedure, building upon the threshold-free network-based statistics, requires setting additional parameters. The PC procedures comprise six variants given the degree of freedom in pooling data, null distribution construction, and controlled error rate. Using numerical simulations, we evaluated the performance of the TT procedure under 600 parameter combinations, then benchmarked TFNOS procedures and baselines across different topologies of effects, sample sizes, and effect sizes, and finally provided illustrative examples with real data. We offer recommended parameter values that allow the TT procedure to stably maintain leading power, while empirically controlling the false discovery rate (FDR) beyond only weakly controlling the familywise error rate (FWER). Notably, the relevant parameters commonly employed in the field appear overly liberal. Furthermore, for the PC procedures, FDR-controlling variants showed improved power compared to FWER-controlling variants, and some of them are simple but do not compromise power. The nonparametric PC procedures allow the selection of any test statistics considered appropriate. Overall, the TFNOS is a generalized framework for inference on edges/nodes of undirected/directed brain networks. We provide empirical and principled criteria for selecting appropriate procedures and may enhance the reproducibility and sensitivity of future brain research.

## 1. Introduction

The brain is a complex network of myriad interacting neurons. Advances in neuroimaging allow us to characterize and investigate brain networks across scales and modalities (Bassett and Sporns, 2017). Node and edge definitions^1^ in brain networks vary across modalities and analysis goals (Bassett and Sporns, 2017; Betzel, 2022). A vast and growing body of research leverages *network neuroscience* tools to study task-related activations, development, nervous system anomalies, and brain–behavior associations, among others (Rubinov and Sporns, 2010; Bassett and Sporns, 2017; Betzel, 2022; Noble et al., 2022; Chai et al., 2023; Hao et al., 2023b). However, brain network studies often involve massive univariate tests on edge- or node-level measures (Helwegen et al., 2023; Hao et al., 2023b). In statistical terminology, they suffer from *multiple testing*/*comparisons problems*. In multiple testing problems, false positives (or type I errors) over all tests must be controlled (Nichols and Hayasaka, 2003; Dmitrienko et al., 2009). In turn, correction procedures often increase false negatives and reduce *statistical power*. Moreover, sample sizes in neuroimaging research are generally small (Button et al., 2013; Yeung, 2018; Szucs and Ioannidis, 2020), with no exception in brain network studies (the median sample size is about 30) (Helwegen et al., 2023). As a result, studies in the field are often underpowered to identify true effects. Underpowered studies also reduce the likelihood that identified effects reflect true effects (Button et al., 2013). Therefore, the development and utilization of statistical procedures that enhance power are paramount for discovering credible findings and increasing the replicability of results.

It only makes practical value to boost power while controlling false positives at an appropriate level. There is no single measure of false positives in studies, e.g., false positive rate (FPR), familywise error rate (FWER), and false discovery rate (FDR)^2^ (Nichols and Hayasaka, 2003; Dmitrienko et al., 2009; Baggio et al., 2018). Common statistical pipelines are primarily of two types: FDR- and FWER-controlling procedures (Noble et al., 2022). Additionally, there are two forms of control over FWER, weak and strong^3^ (Nichols and Hayasaka, 2003; Dmitrienko et al., 2009). Notably, strong control of FWER automatically controls the FDR, but control of the FDR only implies control of the FWER in the weak sense (Nichols and Hayasaka, 2003). Control of FWER weakly often improves the power but has less ability to localize effects (Friston et al., 1996; Nichols and Hayasaka, 2003). When the number of hypotheses is huge, FWER-controlling procedures may become too conservative to detect true effects. Moreover, combinations of different levels of inference with different error controls can generate diverse statistical procedures. Another way may improve the power is to increase the level of inference (Friston et al., 1996; Zalesky et al., 2010; Noble et al., 2022), e.g., from edge-level to cluster-level inference^4^ (Zalesky et al., 2010; Noble et al., 2022). Traditional Bonferroni- and FDR-related procedures (Bland and Altman, 1995; Benjamini and Hochberg, 1995) do not utilize the spatial structure of the data or effects. True effects in brain networks are usually not focused on a single isolated edge or node but are distributed over larger spatial extents. In brain network research, the most popular method for cluster-level inference is the network-based statistic (NBS), which exploits the assumption that edges with effects can form interconnected structures (Zalesky et al., 2010). The NBS is a nonparametric permutation-based method and is an extension of cluster-based inference on voxels in magnetic resonance imaging (MRI) (Bullmore et al., 1999) to brain networks. Put simply, the NBS first performs edge-wise univariate tests. It then thresholds the edge-wise statistics using a preselected threshold and finds connected components (i.e., clusters). Subsequently, it calculates cluster-wise statistics (e.g., the number of edges or the sum of edge-wise statistics in a cluster) and builds an overall null distribution using the largest cluster-level statistic in each permutation. A cluster is declared significant if its cluster-level statistic exceeds the 1 − α quantile of the overall null distribution (Maris and Oostenveld, 2007; Zalesky et al., 2010). Notably, there is no fixed correct option for how to define a cluster, and several variants have been derived from the original NBS (Zalesky et al., 2012; Ge et al., 2021; Zhang et al., 2022).

However, one concerning limitation of the NBS and its variants is the need to preselect a cluster-forming threshold (Zalesky et al., 2010, 2012; Ge et al., 2021; Zhang et al., 2022). This threshold has a large impact on the statistical performance, but no definitive rules guide its choice (Zalesky et al., 2010). It probably creates an unnecessary temptation for researchers and an unquenchable doubt for reviewers. Consequently, threshold-free cluster-level inferences on brain networks have attracted increasing attention and interest (Vinokur et al., 2015; Baggio et al., 2018; Noble and Scheinost, 2020; Vinokur et al., 2023). To date, there are two main ways of realizing them. First, threshold traversal (TT): traverse all possible cluster-forming thresholds and integrate information of clusters identified at all thresholds. One representative approach is threshold-free NBS (TFNBS) (Vinokur et al., 2015; Baggio et al., 2018). Second, predefined clusters (PC): pool data within predefined fixed clusters to calculate cluster-wise statistics, without involving thresholding. One representative approach is constrained NBS (cNBS) (Noble and Scheinost, 2020). The TFNBS is an extension of the threshold-free cluster enhancement (TFCE) method^5^ (Smith and Nichols, 2009) for voxels to brain networks. The cNBS is based on the idea of utilizing predefined brain subnetwork divisions for cluster-level inference (Noble and Scheinost, 2020), consistent with the concept of using atlases to make inferences at the region level instead of the voxel level. Both the TFNBS and cNBS utilize permutation tests to generate null distributions. The difference lies in that the TFNBS constructs an overall null distribution using the maximum statistic (Baggio et al., 2018), whereas the cNBS builds cluster-specific null distributions using cluster-wise statistics (i.e., the average edge-wise statistics within the cluster) (Noble and Scheinost, 2020). This implies that the TFNBS does not require additional multiple-testing corrections, while the cNBS requires corrections for multiple cluster-wise tests. In the present study, we mainly focus on cluster-level threshold-free inferences, as they present a trade-off between the power and capacity for effect localization.

Of note, the field still lacks a threshold-free statistical framework applicable to a wide range of research contexts related to brain networks. Many studies have been devoted to statistical procedures for edges of undirected brain networks (Baggio et al., 2018; Noble and Scheinost, 2020; Noble et al., 2022). Very little attention has been paid to statistical procedures for directed brain networks or nodes. The TT and PC procedures implicitly make assumptions about the spatial structure of the effect of interest (Baggio et al., 2018; Noble and Scheinost, 2020). There is “no such thing as a free lunch”. The contexts in which they are most appropriate for application are different. Moreover, although the TT procedure (e.g., TFNBS) obviates the need to preselect a cluster-forming threshold, it requires setting of two other parameters (i.e., extension and height enhancement parameters: *E* and *H*) (Baggio et al., 2018; Vinokur et al., 2023). Many commonly used toolboxes, e.g., *Matrix3* (Tournier et al., 2019), *CONN* (Nieto-Castanon, 2020), *MNE* (Gramfort et al., 2013), and *FieldTrip* (Oostenveld et al., 2011), have incorporated the TFCE/TFNOS method. However, the recommended values in the TFCE for *t*-statistics of smoothed three-dimensional voxels based on empirical determination and random field theory (Smith and Nichols, 2009) cannot be directly generalized to conventional brain networks and other contexts. The TT procedure based on *F*-statistics may have the potential to confound the effects of different signs (Baggio et al., 2018). When applicable, it appears more appropriate to use the *t*/*z* statistic (Gravetter and Wallnau, 2017; Wooldridge, 2013). More evidence is desperately required to support the choice of *E* and *H* parameters. Furthermore, there is still a lack of comprehensive evaluation across statistical procedures under different spatial structures of effects of interest, effect sizes, and sample sizes. All these factors hinder the selection of appropriate statistical procedures in studies and affect the application of these procedures.

Here, to address these issues, we introduce a statistical framework, named “Threshold-Free Network-Oriented Statistics” (TFNOS), that integrates both the TT and PC procedures. It offers a simple general framework for statistical inferences on edges and nodes in both undirected and directed brain networks. We comprehensively investigate the performance of the TT procedures using numerical simulations with 600 *E*/*H* parameter combinations under three typical effect sizes (Cohen’s *d* = 0.2, 0.5, and 0.8) (Cohen, 1988; Helwegen et al., 2023). The performance metrics include power, FPR, FDR, and FWER. The cluster-level inference maintains strong control of FWER at the cluster level but only weak control of FWER at a finer level (e.g., edge level) (Friston et al., 1996; Zalesky et al., 2010). It would be appealing if suitable *E*/*H* parameters could control the FDR at an appropriate level. Moreover, we comprehensively evaluated the performance across various inference procedures under different spatial extents of effects, and three typical effect sizes (i.e., *d* = 0.2, 0.5, and 0.8), and seven sample sizes (i.e., 10, 20, 50, 100, 200, 500, and 1000 per group) ^6^. As illustration and validation examples, we examined differences between patients with incomplete spinal cord injury (SCI) and healthy controls (HC) in both undirected and directed brain networks. The two main contributions of this study are as follows: (1) We introduce the TFNOS framework that extends from edge-based statistics for undirected brain networks to a unified statistical framework for both nodes and edges of both undirected and directed networks. It considerably expands the application contexts of threshold-free statistical procedures. (2) We provide empirical and principled criteria for the choice of the statistical procedure and parameters for group-level statistical inference in brain networks, contributing to parameter settings of the broadly used toolboxes and stronger conclusions in studies.

## 2. Methods

In this section, we introduce the details of the TFNOS framework (Sections 2.1–2.3) and the experiments (Sections 2.4–2.7) of this study.

### 2.1. Overview of the TFNOS framework

As shown in Fig. 1, the introduced TFNOS framework is a permutation-based framework, which allows for cluster-level statistical inferences on edges and nodes in both undirected and directed networks, especially applicable to brain networks in various modalities and at different scales. This framework comprises two classes of approaches to enable threshold-free inferences: TFNOS-TT and TFNOS-PC. Both the TFNOS-TT and TFNOS-PC performs cluster-level inferences, and the core differences between the two are how to define clusters and utilize cluster-level information. The TFNOS-TT traverses all possible thresholds on the edge-/node-wise statistic map and integrates information from clusters (data-driven) identified from suprathreshold edges/nodes at all thresholds. The TFNOS-PC pools data within predefined clusters to obtain cluster-wise statistics without involving a thresholding process. Details of the TFNOS-TT and TFNOS-PC are described in Sections 2.2– 2.3. The TFNOS framework provides multiple cluster definition options depending on the application context and the topology of the effect of interest. It is a generalized framework for group-level statistical inference in network neuroscience.

**Fig. 1.**
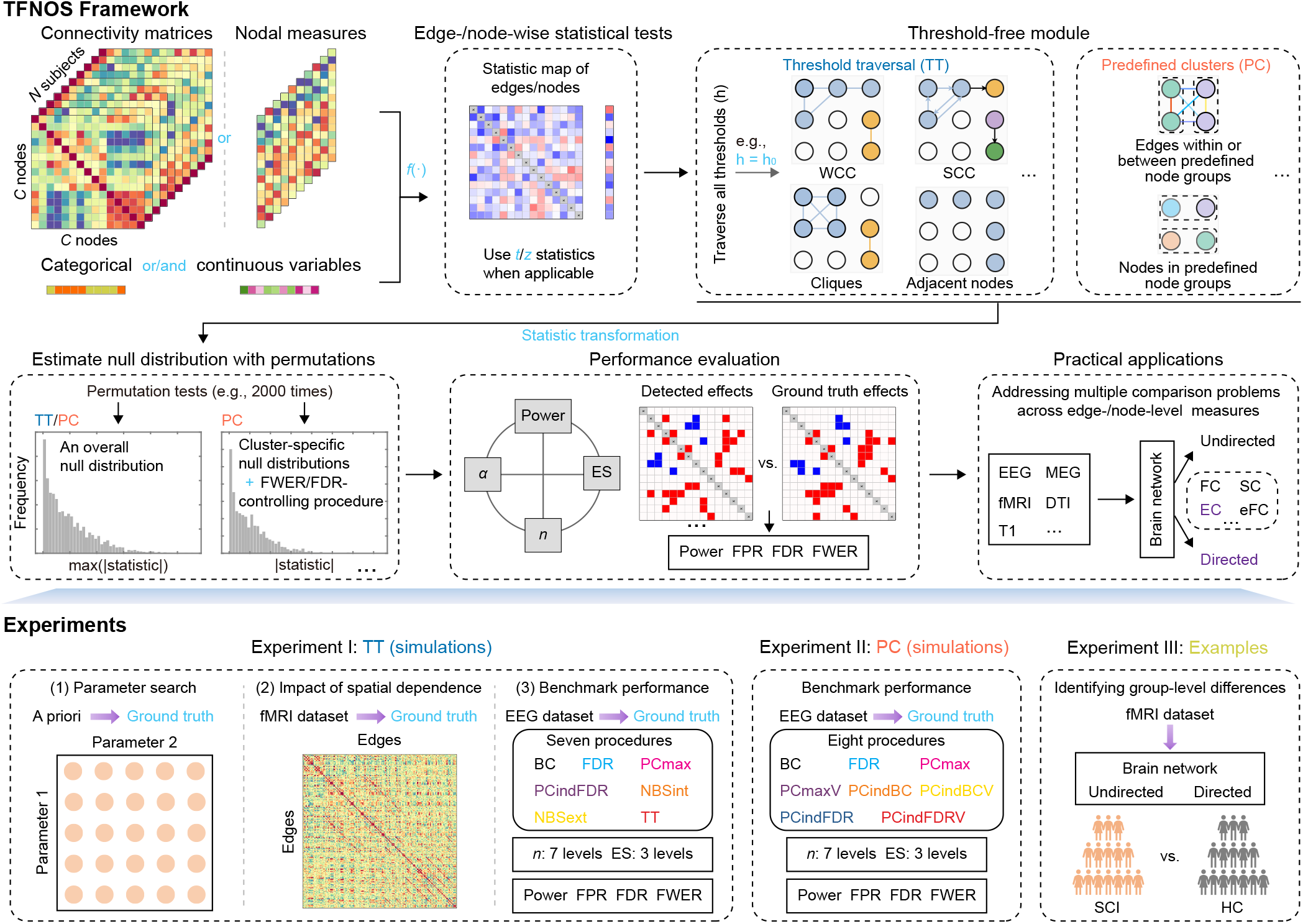
Overview of TFNOS framework and experiments in the present study. (Top) Schematic illustration of TFNOS framework. The TFNOS is a permutation-based framework for cluster-level statistical inference in undirected and directed brain networks. The input “independent” variable can be the connectivity matrix or node-level measures. The input “dependent” variable can be a group indicator variable or continuous variable, etc. This framework has many degrees of freedom in terms of the way of defining clusters, the way of calculating cluster-level statistics, the form of null distribution construction, and which error rate to control. See Sections 2.1–2.3 for more details of the TFNOS framework. (Bottom) Experiments in this study. Using numerical simulation, we benchmark the performance of the TT procedure and baselines, and the PC procedures (six variants) and baselines across seven sample sizes and three effect sizes. See Sections 2.5.3 and 2.6 for explanations of code names of the statistical procedures. As illustrations, the TFNOS framework is applied to a real fMRI dataset. See Sections 2.4– 2.7 for more details of the experiments. TFNOS = threshold-free network-oriented statistics; WCC = weakly connected component; SCC = strongly connected component; α = significant level; *n* = sample size; ES = effect size; FPR = false positive rate; FDR = false discovery rate; FWER = familywise error rate; EEG = electroencephalography; MEG = magnetoencephalography; fMRI = functional magnetic resonance imaging; DTI = diffusion tensor imaging; T1 = T1 weighted images; FC = functional connectivity; SC = structural connectivity; EC = effective connectivity; eFC = edge-centric functional connectivity. SCI = patients with incomplete spinal cord injury; HC = healthy controls.

### 2.2. TFNOS-TT

The TFNOS-TT is inspired by and extends the previous TFCE/TFNBS (Smith and Nichols, 2009; Baggio et al., 2018). The TFCE method was first proposed in the work of Smith and Nichols (2009) for voxel-based analysis and was translated to edge-based analysis in undirected networks in the work of Vinokur et al. (2015) and Baggio et al. (2018).

Here, we expand upon the existing statistical procedure (Vinokur et al., 2015; Baggio et al., 2018) to encompass group-level inferences on nodes and edges in both undirected and directed brain networks across scales and modalities. The key to realizing the extensions is how to define clusters. The true effect size is generally unknown (Lakens, 2022), and the cluster definitions need to consider the spatial topologies of effects of interest. For instance, the consequences of focal brain damage extend far beyond the lesion site, propagating to subnetworks formed by the affected area and other brain regions that are connected to it structurally or functionally (Bartolomeo and Thiebaut de Schotten, 2016; Siegel et al., 2016). In this case, it is plausible to define clusters by leveraging interconnected subnetworks influenced by the effects (Zalesky et al., 2010; Baggio et al., 2018). We introduce four kinds of cluster definitions for the TFNOS-TT, and illustrative examples are presented in Fig. 1.

- A cluster can be defined as the edges within a weakly connected component (WCC). A WCC is a subnetwork of a network, e.g., functional connectivity (FC); effective connectivity (EC), structural connectivity (SC), and edge-centric FC (eFC) (Faskowitz et al., 2020; Hao et al., 2023b), where all nodes are connected to each other by some edges, ignoring the direction.
- A cluster can be defined as the edges within a strongly connected component (SCC). An SCC is a subnetwork of a directed network, e.g., EC, where every node can reach the other node by the directed path.
- A cluster can be defined as the edges within a maximal clique derived. A clique is a subset of nodes of an undirected network such that every node is connected to the other nodes in the clique by an edge. A clique is said to be a maximal clique if it is not a proper subset of any other clique.
- A cluster can be defined as spatially adjacent nodes. Whether two nodes are neighboring or not can also be exploited simultaneously with their functional and structural proximity (Mansour Lakouraj et al., 2022).

We briefly introduce how the TFNOS-TT realizes “threshold-free” in the context of undirected brain networks (e.g., with *C* brain regions/nodes). Univariate statistics of interest are performed across all edges to obtain edge-wise test statistics (*C* × *C*). The TFCE converts the original edge-wise test statistics into “threshold-free” edge-wise cluster-based scores ***Q***^*C*×*C*^ (referred to as TFCE scores), following Eq. (1).

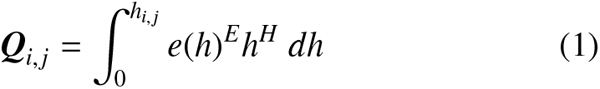

where ***Q***_*i*, *j*_ denotes the transformed score of edge {*i, j*} between nodes *i* and *j, h*_*i*, *j*_ is the original test statistic of this edge, *h* is the current threshold, *e*(*h*) is the size of the cluster containing this edge at threshold *h*, and *E* and *H* are the extension and height enhancement parameters, respectively. In practice, this integral is numerically approximated by a finite sum using the thresholding interval *dh*. A cluster can be defined as a set of edges within a WCC (Baggio et al., 2018) or clique (Zhang et al., 2022) detected from the suprathreshold network (Zalesky et al., 2010; Baggio et al., 2018). The cluster size is defined as the number of edges in the cluster.

For the TFONS-TT procedure, we recommend using edge-wise *t*/*z* statistics as input, whenever possible, for three reasons. (1) They can separate the effects of different signs, which is crucial as the composition of edges with different effect signs in a cluster may violate the implicit assumptions regarding clusters. (2) They can be obtained easily from many types of hypothesis testing. For example, the *t*-statistic can be directly obtained in paired or two-sample *t*-tests. The *z*-statistic can be obtained in Mann-Whitney or Wilcoxon tests by a normal approximation when the sample size *N* is relatively large (e.g., *N* > 20). The correlation coefficient of Pearson or Spearman correlation can be converted to the *t*-statistic (Gravetter and Wallnau, 2017). (3) In generalized linear regression, when we are interested in whether a single coefficient (e.g., of the dummy variable for the group) is different from zero, the corresponding *t*-statistic is best suited for this case but the *F*-statistic is to detect whether a set of coefficients is different from zero (Wooldridge, 2013).

The algorithms for the TFNOS-TT statistical inference on edges (Fig. 2) and nodes (Fig. S1) in brain networks are outlined through pseudocodes. For the sake of completeness of the content, the four main steps of the algorithm are presented using the example of investigating differences between two groups of subjects in FC:

**Fig. 2.**
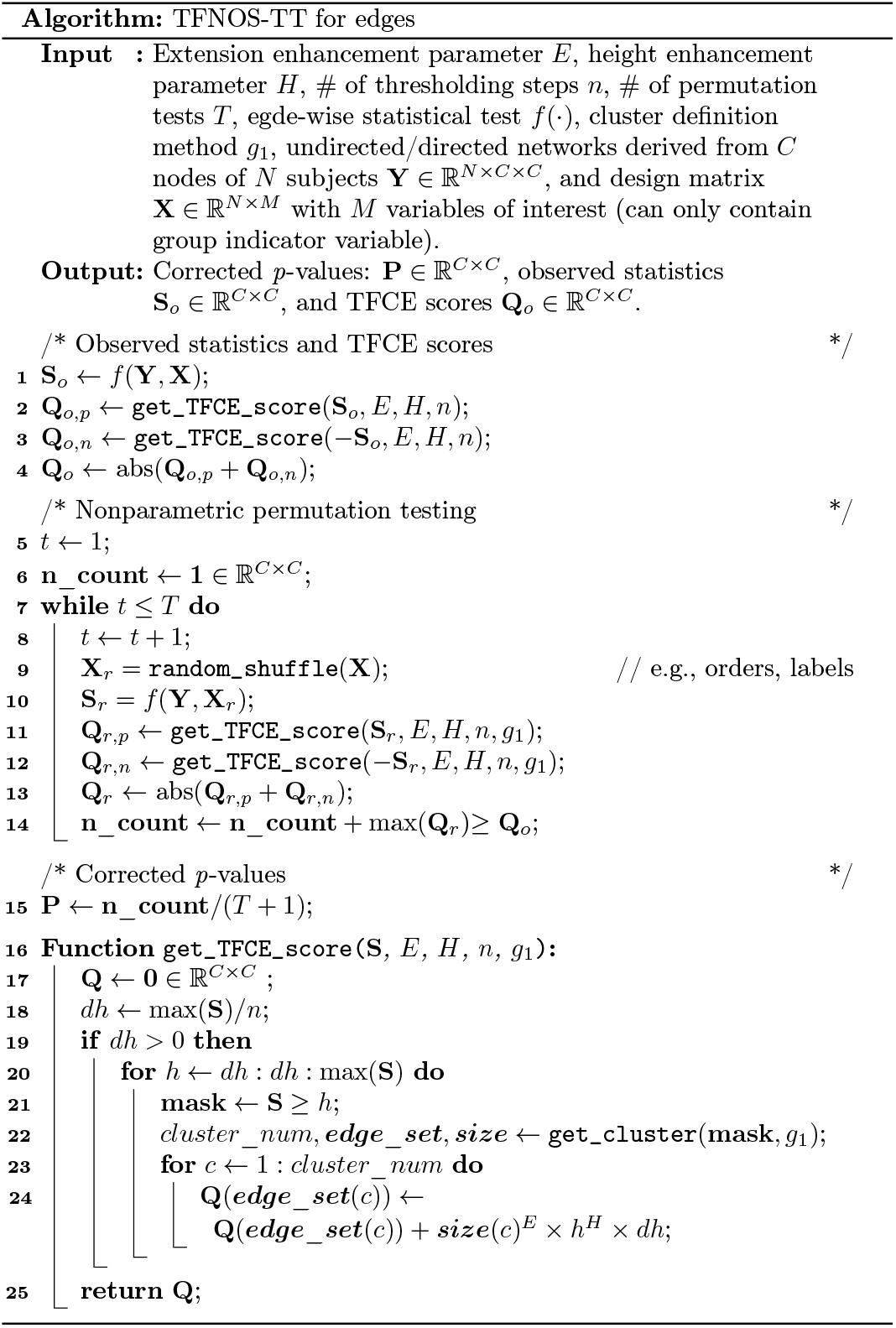
Pseudocodes of the TFNOS-TT procedure for statistical inference on network edges. See Fig. S1 for pseudocodes of the TFNOS-TT procedure for network nodes.

1. Calculate the observed edge-wise test statistics (e.g., *t*-statistic) and the TFCE transformed scores for both positive and negative effects, if applicable.
2. Randomly shuffle the group labels and calculate the edge-wise statistics and TFCE transformed scores on this randomized partition, recording the maximum absolute score across all edges.
3. Repeat step (2) a large number of times (e.g., 2000 times)^7^.
4. The corrected *p*-value (Monte Carlo *p*-value) of one edge is determined as the proportion of random partitions that the maximum absolute score is larger than the observed absolute statistic of this edge.

To preserve the spatial structure, the permutation is not performed independently edge by edge. Instead, the entire network is shuffled as a whole. The algorithm for edges differs from that for nodes, where an additional nodal relationship matrix is required to describe the adjacency of each node to other nodes.

### 2.3. TFNOS-PC

The TFNOS-PC takes into account the fact that in some contexts, such as task-related activities (Noble et al., 2022), and severe brain injury (Hao et al., 2023a), the underlying effects are widespread, involving a large number of edges in the brain network. In contrast to the cluster definitions described in TFNOS-TT, the clusters in TFNOS-PC are all pre-defined (Fig. 1). Specifically, with regard to nodes, clusters can be defined as predefined node groups, which are determined by community detection (Garcia et al., 2018; Wierzbiński et al., 2023) or clustering (Thomas Yeo et al., 2011; Yan et al., 2023) algorithms, or even customized by the researcher based on empirical knowledge. For edges, clusters can be defined as the set of edges within or between predefined node groups. In extreme cases, each edge or node may be regarded as a cluster, and the inferences are at the edge level.

Here, we introduced TFNOS-PC procedures with different variants (see also Section 2.6) given the degree of freedom in pooling data, null distribution construction, and controlled error rate. The algorithms of TFNOS-PC statistical inference on network edges (Fig. 3) and nodes (Fig. S2) are presented by the pseudocode. The four primary steps of the TFNOS-PC algorithm are illustrated using an example that investigates differences in FC at the (predefined) cluster level between two groups of subjects:

**Fig. 3.**
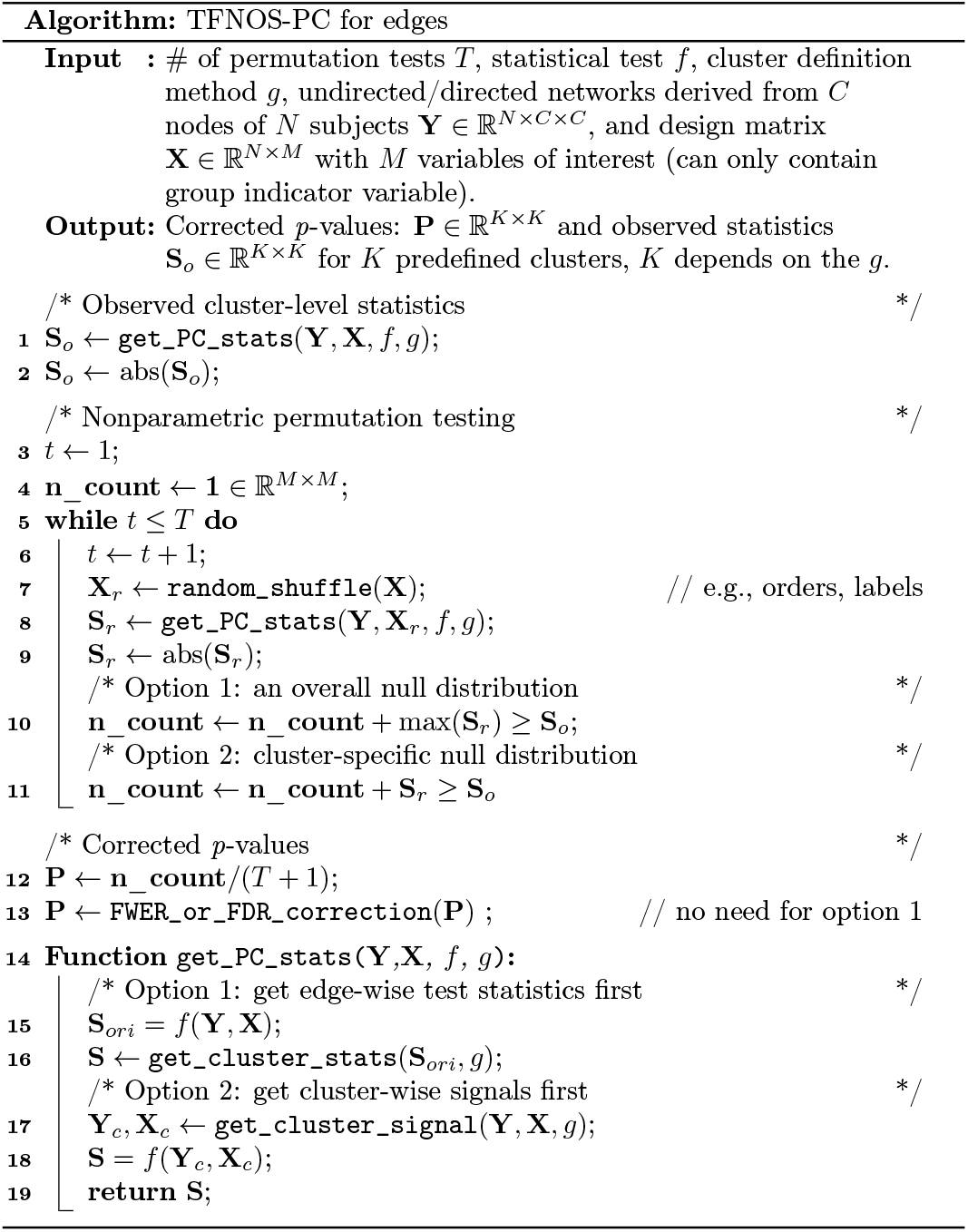
Pseudocodes of TFNOS-PC procedures for statistical inference on network edges. See Fig. S2 for pseudocodes of TFNOS-PC procedures for network nodes.

1. Calculate the observed cluster-level test statistics. There are two options. One is to perform the edge-wise statistical test within the cluster and then average the edge-wise statistics across all edges (Noble and Scheinost, 2020). The other is to calculate the mean edge value within the cluster and subsequently perform the tests to get cluster-wise statistics.
2. Randomly shuffle the group labels then compute the cluster-wise statistic on this random partition. The maximum absolute cluster-wise statistic across all clusters (option 1) or all absolute cluster-wise statistics (option 2) is then recorded.
3. Repeat step (2) a large number of times (e.g., 2000 times).
4. The Monte Carlo *p*-value of a cluster is calculated as the proportion of random partitions that the maximum absolute statistic is larger than the observed statistic of this cluster for option 1 in step (2) or the proportion of random partitions that the absolute statistic of this cluster is larger than the observed absolute statistic of this cluster for option 2 in step (2). If option 2 is used in step (2), the *p*-values should corrected for the multiple tests (28 tests in this example), e.g., Bonferroni or FDR correction.

As noted above, there is considerable flexibility in the implementation of the TFNOS-PC inference. We pursue simple and powerful statistical procedures and experimentally compare the performance of different variants. Of note, there exist hybrid methods. For example, the semi-cNBS (scNBS) method proposed in (Dai et al., 2022), which is based on the cNBS, but utilizes additional data to select an “optimal” threshold for further selection of the elements within the clusters. In general, the hybrid methods either require larger sample sizes or stronger a priori of effect distribution. Evaluations of these hybrid methods are excluded from the present study, although they may achieve good performance in certain scenarios.

### 2.4. Overview of the experiments

The current study comprises three main experimental components (Fig. 1). In Experiments I and II, simulation experiments were conducted to evaluate the performance of TFNOS and traditional procedures (e.g., Bonferroni, FDR, and NBS). Additionally, Experiment I involved an investigation of the appropriate parameter (*E* and *H*) range for TFNOS-TT and an examination of the impact of the data spatial dependence on the statistical performance. In Experiment III, the TFNOS framework was applied to a real clinical dataset to detect group-level differences in both FC and EC, serving as illustrative examples. A detailed description of the experiments is expanded in the following Sections 2.5–2.7. Of note, the experiments are mainly in the context of statistical inference on the edges. Statistical tests on edges suffer from a more serious multiple-testing problem than nodes. Here, we provide an outline of the ground truth setup, simulated data generation, and estimation of performance metrics.

#### Ground truth setup and real datasets

To give some realistic basis for the topology of the ground-truth effect, we refer to the structure of significant clusters reported in previous studies (Ge et al., 2022; Chen et al., 2023) or leverage real datasets to generate the ground-truth effects similar to the approach used in Baggio et al. (2018). The real datasets include a functional MRI (fMRI) dataset and an EEG dataset. The fMRI dataset collected data from 48 patients with SCI and 29 HCs. The EEG dataset contains source-reconstructed EEG data from 178 patients with prolonged disorders of consciousness. Data acquisition and preprocessing of the fMRI and EEG data can be found in Ge et al. (2021) and Hao et al. (2023a), respectively. This study was approved by the Ethics Committee of Beijing Tsinghua Changgung Hospital, the Ethics Committee of the PLA Army General Hospital, and the Institutional Review Board of Tsinghua University. Written informed consent was obtained from each participant or their legal surrogates.

#### Simulation data generation

In the present study, all simulation experiments were conducted in the context of between-subjects designs and followed similar data generation rules, generating data for “healthy subjects” and “patients” (Baggio et al., 2018). For each “healthy subject”, the values of each edge are randomly sampled from a normal distribution 𝒩 (0, 1). For each “patient”, the values of edges with a true effect were randomly sampled from a normal distribution N (*d*, 1)^8^, and those of the other edges were randomly sampled from a normal distribution 𝒩 (0, 1). For both “healthy subjects” and “patients”, a symmetric matrix (undirected) of dimension *C* × *C* (subject to the experiment setup) is generated with all diagonal elements 1. We performed simulations for three typical effect sizes with *d* of 0.2 (small), 0.5 (medium), and 0.8 (large) (Cohen, 1988; Helwegen et al., 2023). Considering the time consumption of the experiments, for each level of effect size, 100 simulation data groups were generated. Each data group comprises *n* “healthy subjects” and *n* “patients”. For permutation-based procedures, 2000 times permutations were performed, and the *t*-statistics (two-sample *t*-tests) were utilized.

#### Estimation of performance metrics

Four metrics, including power, FPR, FDR, and FWER, are utilized to evaluate the statistical performance. For NBS and TFNOS-TT inferences, it is important to note that clusters identified by algorithms from the current data do not necessarily have to completely align with the clusters formed by ground-truth effects. In the TFNOS-TT, the TFCE score represents an “edgelization” of clusters, where edge-level statistics are computed by “borrowing” statistics from all edges in the cluster under all possible cluster-forming thresholds (Smith and Nichols, 2009; Baggio et al., 2018). Consequently, explicit calculation of these metrics at the cluster level is precluded, leading to an intuitive idea of computing them at the edge level. This requires assuming that the null is false for any individual edge in a cluster if the null for this cluster is rejected. Theoretically, for NBS, declaring a cluster to be significant only implies that there is an effect on at least one edge in this cluster (Noble et al., 2022). For TFNOS-TT inferences, rejecting the null at an edge implies that there exists at least one cluster containing that edge, with an effect on one or more edges in the cluster (Smith and Nichols, 2009; Noble et al., 2022). However, in simulation experiments where the precise location of the true effect is known, it is possible to investigate whether the presence of appropriate *E* and *H* parameters allows for the interpretation of results at the edge level (e.g., empirical control of type I errors). Others have adopted a similar practice: cluster-level inference to calculate performance metrics at the edge level (Noble et al., 2022; Baggio et al., 2018; Zalesky et al., 2010). Furthermore, for TFNOS-PC procedures, the matrices were computed at the same spatial scale as the level of inference (i.e., cluster level). The test is two-tailed and the significant level is 0.05 in all experiments.

### 2.5. Experiment I: TFNOS-TT simulations

#### 2.5.1. Parameter selection

In the TFNOS-TT, the selection of a hard cluster-forming threshold (Zalesky et al., 2010; Maris and Oostenveld, 2007) is avoided, but the *E* and *H* parameters need to be chosen (see Section 2.2). Previous research has suggested recommended values of *E* and *h* in certain contexts (Smith and Nichols, 2009; Baggio et al., 2018), but it is uncertain whether these recommended values are suitable for various experimental designs involving brain networks. For example, the recommended values in Smith and Nichols (2009) are based on the context of smoothed three-dimensional voxel-based data (input raw *t*/*z* statistic matrix), while those in Baggio et al. (2018) are obtained in the context of brain networks using the *F*-statistic matrix as the input. According to Eq. (1), and the choice of test statistics can impact the TFCE score. See also Section 2.2 for the rationale behind recommending the use of the *t*/*z* statistic (if applicable).

*Search space*. We conducted evaluations for the TFNOS-TT procedure on a range of 30 *E* (0.05–1.50 at intervals of 0.05) and 20 *H* (0.5–10.0 at intervals of 0.5) parameter values, in a total of 600 *E*/*H* parameter combinations. The searched space encompassed all previously recommended *E*/*H* values (Smith and Nichols, 2009; Baggio et al., 2018; Tournier et al., 2019).

##### Ground truth setup

In simulation experiments, the precise location and topology of the ground-truth effects need to be set in advance. Here, we partially referred to the topologies of significant clusters reported in prior case-control studies (Ge et al., 2022; Chen et al., 2023) and designed hybrid topologies by combining topologies such as star, ring, and tree. As depicted in Fig. S3, the hybrid topologies consist of five connected components with a total of 91 edges. From largest to smallest, these five connected components contain 51 edges (41 nodes), 35 edges (28 nodes), 5 edges (6 nodes), 3 edges (4 nodes), and 1 edge (2 nodes), respectively. Among these, the largest, fourth, and fifth were set with negative effects (i.e., “patients” < “healthy subjects”), and the second and third were set with positive effects (i.e., “patients” > “healthy subjects”).

##### Simulations

For each *E*/*H* parameter combination from the search space (600 *E*/*H* parameter combinations) and each effect size level (i.e., 0.2, 0.5, and 0.8), 100 simulation data groups were generated. Each data group comprises 200 “healthy subjects” and 200 “patients”. The cluster used here is defined as the edges within a WCC. Additionally, 100 simulation data groups were generated with each group also composed of 200 “healthy subjects” and 200 “patients”, but all values of the simulation data groups were randomly sampled from a normal distribution 𝒩 (0, 1). These data groups were used to estimate weak-sense FWER.

#### 2.5.2. Impact of spatial dependence

Prior research on brain network simulations has exploited the topology of the effects of interest but has not considered the spatial dependence of the data (edges) (Zalesky et al., 2010; Baggio et al., 2018; Zhang et al., 2022). Here, we utilized a method based on Cholesky decomposition to evaluate the impact of spatial dependency between edges on the statistical performance with a toy example. More specifically, the Pearson correlation coefficient was used to measure the dependence between edges. Let ***R*** ∈ ℝ^*M*×*M*^ be the Pearson correlation matrix of *M* edges, with *r*_*i*, *j*_ denoting the linear dependency strength between edges *i* and *j*. If ***R*** is positive definite, it can be decomposed by Cholesky decomposition as ***R*** = ***U***^*T*^ ***U***, where ***U*** ∈ ℝ^*M*×*M*^ is an upper triangular matrix, and the 2-norm of column vectors of ***U*** is 1. Let *X*_1_, *X*_2_, …, *X*_*M*_ be i.i.d. normal random variables with means 0 and variances 1, and then the linear combination 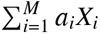 follows a normal distribution 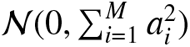. Correlated normal variables [*Z*_1_, *Z*_2_, …, *Z*_*M*_] can be obtained by multiplying the uncorrelated normal variables [*X*_1_, *X*_2_, …, *X*_*M*_] with ***U***, and they still follow a standard normal distribution. The correlation between *Z*_*i*_ and *Z*_*j*_ will be *r*_*i*, *j*_ as expected.

##### Ground-truth setup

The fMRI dataset was utilized to generate the matrix ***R*** and set the ground-truth edges. Specifically, the automated anatomical labeling (AAL) atlas (Tzourio-Mazoyer et al., 2002) was used to parcellate the brain into 116 brain regions, and the Pearson-correlation-based FC was computed between each pair of region-level average signals. Let ***X*** ∈ ℝ^*N*×*M*^ be a matrix consisting of the edge values of *N* subjects and *M* edges for each subject (column-wise *z*-scored), and *rank*(***X***) = *min*(*N, M*). The correlation/covariance matrix ***R*** ∈ ℝ^*M*×*M*^ (i.e., ***X***^*T*^ ***X***) is positive semi-definite, since 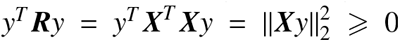, ∀*y* ≠ 0. If *M* ⩽ *N*, then the rank of ***X*** and ***R*** is *M*, and thus ***R*** is a full rank and positive definite matrix, allowing for Cholesky decomposition. The edge-wise difference between the SCI and HC groups was examined by two-sample *t*-tests. Given that the sample size of the SCI and HC groups is 48 and 29, respectively, a subnetwork with 8 nodes (8 ×7/2 = 28 edges) that satisfies *M* ⩽ *N* for both groups^9^ was selected and a threshold (observed *d* = 0.8) was applied to the *t*-statistic matrix of this subnetwork, defining the suprathreshold edges (*n* = 7) as group truth (see Fig. S4).

##### Simulations

To investigate whether dependence notably influences statistical performance in this toy example, we conducted simulation experiments with three effect sizes (i.e., 0.2, 0.5, and 0.8) and seven sample sizes (i.e., 10, 20, 50, 100, 200, 500, and 1000 per group). According to the results from Section 2.5.1 (see Section 3.1 for details), the parameters *E* and *H* were set to 0.4 and 3, respectively.

#### 2.5.3. TFNOS-TT performance evaluation

In this section, we include the TFNOS-TT procedure in an empirical benchmark with other six statistical procedures.

The simulation experiments were conducted with the between-group design in the context of FC, using two-sample *t*-tests for edge-wise tests. The seven procedures were assigned name codes and briefly described as follows:

1. BC: Bonferroni correction procedure (Bland and Altman, 1995);
2. FDR: FDR correction procedure (Benjamini and Hochberg, 1995);
3. PCmax: TFNOS-PC-based procedure (each edge is regarded as a cluster) with an overall null distribution by the maximum absolute statistic across all edges;
4. PCindFDR: TFNOS-PC-based procedure (each edge is regarded as a cluster) with edge-wise null distribution and subsequent FDR correction for multiple tests;
5. NBSint: NBS procedure with the cluster size defined as the sum of the edge-wise statistics within the cluster (WCC) (Zalesky et al., 2010);
6. NBSext: NBS procedure with the cluster size defined as the number of edges in the cluster (WCC) (Zalesky et al., 2010);
7. TT: TFNOS-TT-based procedure with the cluster defined as the set of edges in a WCC;

##### Ground-truth setup

We configured the edges with the group-truth effect using the EEG dataset. The “Yan2023” homotopic 100-area cortex atlas (Yan et al., 2023) was employed to extract EEG source-space region-level time series and region-to-region FC was computed using the weighted phase-lag index (Vinck et al., 2011) (see Hao et al. (2023a) for further details). The edge-wise Mann–Whitney *U* test was performed to evaluate the difference between patients in the vegetative state/unresponsive wakefulness syndrome state (VS/UWS) and in the minimally conscious state (MCS). The suprathreshold edges, obtained by thresholding the absolute values of *z*-statistic matrix by 3, were set with the groud-truth effect. As depicted in Fig. S5, they are comprised of two WCCs: one with 69 edges (67 nodes) and the other with 1 edge (2 nodes).

##### Simulations

For all seven statistical procedures, no hard threshold is required except for NBSint and NBext, for which the threshold was set to |*t*| = 3 in line with the value used in the NBS method proposed study (Zalesky et al., 2010). For the TT procedure, the parameters *E* and *H* were set as 0.4 and 3, respectively. Additionally, simulation experiments were conducted with three different effect sizes and seven different sample sizes.

### 2.6. Experiment II: TFNOS-PC simulations

In this section, We benchmarked the TFNOS-PC procedures and two other traditional procedures using simulation experiments. The TFNOS-PC procedures consist of six variants that differ in the construction of the null distribution, calculation of cluster-wise statistics, and error control type (see Section 2.3). The eight procedures are name-coded and briefly described as follows:

1. BC: Cluster-wise tests on cluster-level average signals + Bonferroni correction procedure (Bland and Altman, 1995);
2. FDR: Cluster-wise tests on cluster-level average signals + FDR correction procedure (Benjamini and Hochberg, 1995);
3. PCmax: Cluster-wise statistics derived from edge-wise statistics within the clusters + An overall null distribution using the maximum absolute statistic across all cluster-wise statistics (variant 1);
4. PCmaxV: Cluster-wise statistics derived from cluster-level average signals + An overall null distribution using the maximum absolute statistic across all cluster-wise statistics (variant 2);
5. PCindBC: Cluster-wise statistics derived from edge-wise statistics within the clusters + Cluster-specific null distributions + Bonferroni correction procedure (Noble and Scheinost, 2020) (variant 3);
6. PCindBCV: Cluster-wise statistics derived from cluster-level average signals + Cluster-specific null distribution + Bonferroni correction procedure (variant 4);
7. PCindFDR: Cluster-wise statistics derived from edge-wise statistics within the clusters + Cluster-specific null distributions + FDR correction procedure (Noble and Scheinost, 2020) (variant 5);
8. PCindFDRV: Cluster-wise statistics derived from cluster-level average signals + Cluster-specific null distributions + FDR correction procedure (variant 6).

#### Ground truth setup

We utilized the EEG dataset to generate the cluster-level group truth. The 100-area “Yan2023” atlas (Yan et al., 2023) can be matched to the “Yeo2011” seven resting-state networks (RSNs) (Thomas Yeo et al., 2011), which were considered as the predefined node groups for the present study. The average connectivity between or within RSNs was computed for each recording. Then, the group differences between patients in MCS and in VS/UWS were tested using the Mann–Whitney *U* test. A threshold of| *z* |= 2.0 was applied and 9 clusters (2 within RSNs and 7 between RSNs) out of 28 clusters (7 within RSNs and 21 between RSNs) were set with the groud-truth effect, as shown in Fig. S6.

#### Simulations

We conducted simulation experiments with three different effect sizes and seven different sample sizes. All simulated edge values were sampled from the corresponding normal distribution (see Section 2.4).

### 2.7. Experiment III: Test in the fMRI dataset

In this section, as illustrative examples, we applied the TFNOS-TT procedure and TFNOS-PC (PCindFDRV) to the fMRI dataset to explore brain network alterations in patients with SCI (*n* = 48) compared to HC (*n* = 29). The sample sizes for both groups are representative of typical sample sizes in case-control brain network studies (Helwegen et al., 2023). In addition to constructing undirected brain networks based on Pearson correlations (Pearson-FC), we also used the approach of convergent cross-mapping (CCM) to generate directed brain networks (CCM-EC) (Sugihara et al., 2012; Ye et al., 2015; Ge et al., 2021). The linear correlation cannot reflect the nonlinear coupling between signals, and it is difficult to have a causal interpretation, while the CCM method is suitable for detecting causal relations between the signals in a not entirely random system (i.e., underlying manifold governing the dynamics) (Sugihara et al., 2012; Yuan and Shou, 2022). The brain was parcellated into 116 regions using the AAL atlas. For the Pearson-FC and CCM-EC, the dimension of the connectivity matrix is 116 × 116.

For the TFNOS-TT procedure, a cluster is defined as the set of edges within each WCC for the Pearson-FC and within each SCC for the CCM-EC. The parameters *E* and *H* were set as 0.4 and 3.0, respectively. For the TFNOS-PC procedure, the 116 regions were divided into 14 node groups (7 left and 7 right) based on their anatomical locations (frontal, limbic, temporal, basal ganglia, parietal, occipital, and cerebellum). There are 105 predefined clusters (14 within node group and 14 × 13/2 = 91 between node groups) for P-FC and 196 predefined clusters (14 within node group and 14 × 13 = 182 between node groups) for CCM-FC.

## 3. Results

### 3.1. Experiment I: TFNOS-TT simulations

#### 3.1.1. Existence of appropriate E and H parameters

We first evaluated the impact of *E* and *H* parameters (600 *E*/*H* combinations) on statistical performance using numerical simulations. The topology of the ground truth effect can be found in Fig. S3. For each of the three effect sizes (i.e., 0.2, 0.5, and 0.8), we generated 100 simulation datasets and computed empirical performance metrics for each *E*/*H* parameter combination.

Fig. 4 shows the power, FPR, and FDR of the TFNOS-TT procedure under each of the 600 *E*/*H* parameter combinations, with effect sizes of 0.2, 0.5, and 0.8. See Fig. S7 for the results of weak- and strong-sense FWER. We observed that the performance metrics are sensitive to the choice of *E*/*H* parameters. Broadly speaking, when *E* is less than 1.0, higher *E* and lower *H* lead to increased power but also elevated FPR and FDR. Moreover, as expected, the TFNOS-TT procedure controls the FWER only in the weak sense at the edge level (Fig. S7). Nevertheless, there exist appropriate *E* and *H* parameters (not unique) that trade off power and type I error rate, and interestingly, FDR can be empirically controlled to a low level (e.g., < 10%) (Fig. 4). We recommend a value of 0.4 for *E* and 3.0–7.0 for *H*, with larger values of *H* being more conservative. Under these values, the results show that power remains at a relatively high level with a relatively low FDR (e.g., *H* = 3.0, FDR < 10%; *H* = 7.0, FDR < 5%). Notably, these recommended values of *E* and *H* are in the context of inferences on edges (*t*-statistics). We provided an example of *E*/*H* parameter search in the context of inferences on nodes (EEG channels) in the Supplementary Material (Figs. S8–S10), where *E* = 1.0 and *H* = 2.0–3.0 are appropriate.

**Fig. 4.**
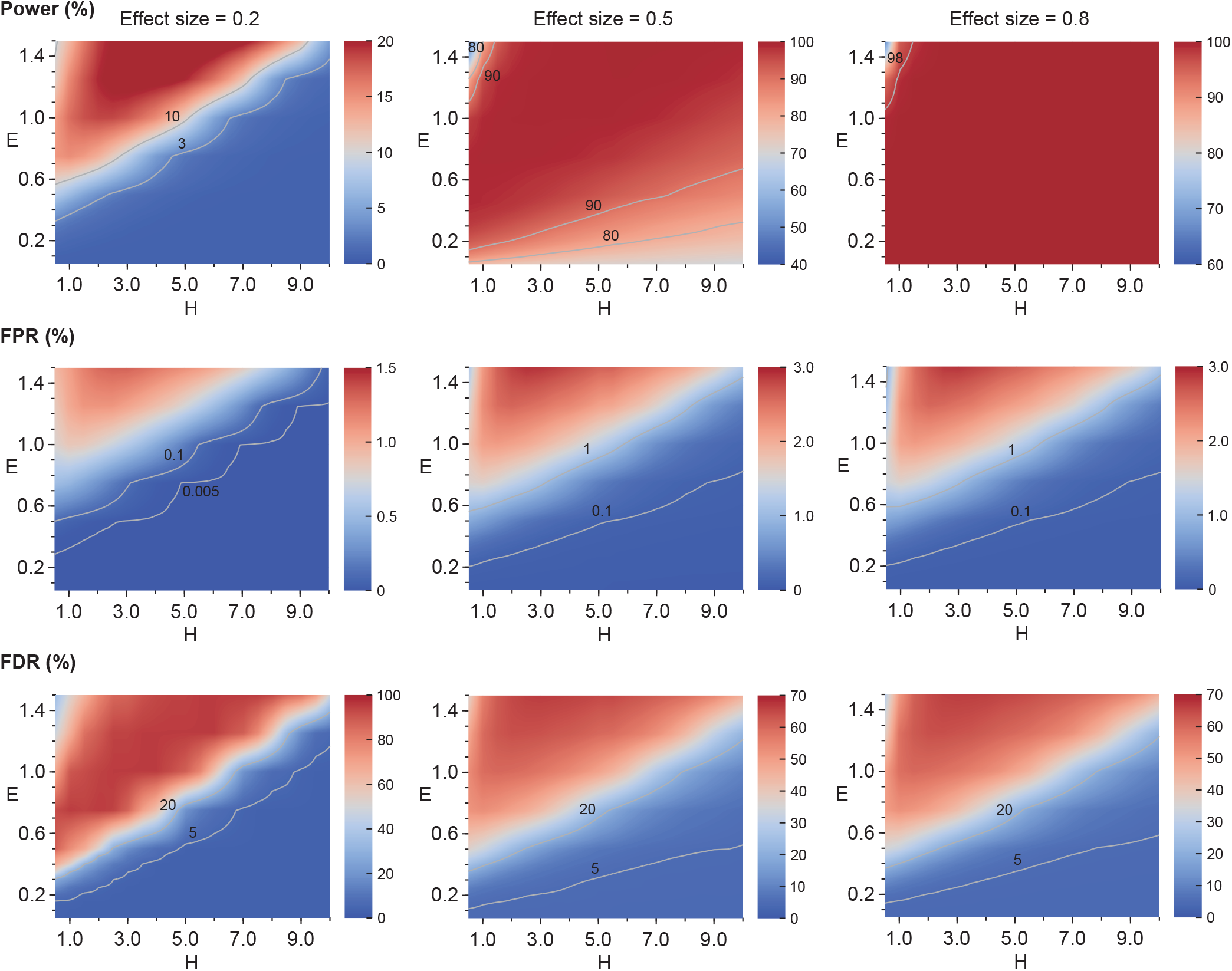
Parameter search of TFNOS-TT procedure. Heatmaps display empirical power (top), FPR (middle), and FDR (bottom) under numerical simulations (100 repetitions) for 600 *E*/*H* parameter combinations (30 *E* and 20 *H* values), with effect size (Cohen’s *d*) of 0.2 (left), 0.5 (middle), and 0.8 (right). For better visualization, the data for the heatmaps were linearly interpolated somewhat, and some necessary contours were added (in gray). See Fig. S7 for the heatmaps of FWER.

#### 3.1.2. Impact of spatial dependence

In addition to the effect topology, we investigated the potential influence of spatial dependence on statistical performance (see Section 2.5.2). In this experiment, *E* and *H* were set to 0.4 and 3.0, respectively. Fig. 5 illustrates the power, FPR, FDR, and strong-sense FWER of the TFNOS-TT procedure with considering and not considering spatial dependence. The results from this experimental scenario show that the impact of simulated data, with or without considering spatial correlation, on statistical performance is slight, particularly for power. In the provided toy example (Fig. S4), the FDR was also empirically controlled across three effect sizes and seven sample sizes (≤ 5%). Additionally, no consistent difference pattern was observed in FPR and strong-sense FWER. As expected, cluster-level inferences cannot control FWER in the strong sense at the edge level. The preliminary findings require the reader to note that it remains to be determined the generalizability of these results to a widespread of scenarios beyond the examined data.

**Fig. 5.**
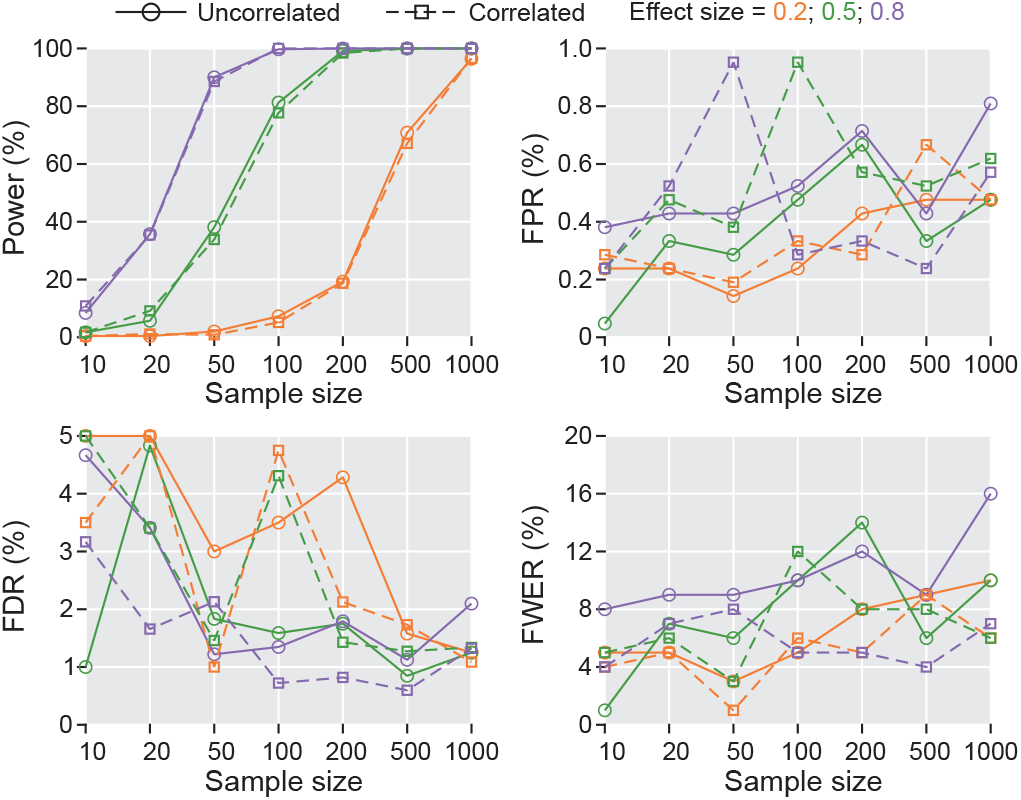
Impact of spatial dependence among edges. The line plots show the power, FPR, FDR, and strong-sense FWER of the TFNOS-TT procedure at different combinations of effect sizes and sample sizes (per group) with (dashed lines) and without (solid lines) considering spatial correlation.

#### 3.1.3. Benchmarking performance of the TFNOS-TT procedure and baselines

We conducted an empirical benchmark of the performance of the TT and six other procedures using simulated datasets. Notably, the performance metrics of all seven procedures were calculated at the edge level (see Section 2.4). Moreover, it is notable whether the TT procedure can demonstrate control of the FDR in this new context under the *E*/*H* parameters recommended in Section 3.1.1.

As shown in Fig. 6, for a given statistical procedure, its power increases with the rise in sample size or effect size.

**Fig. 6.**
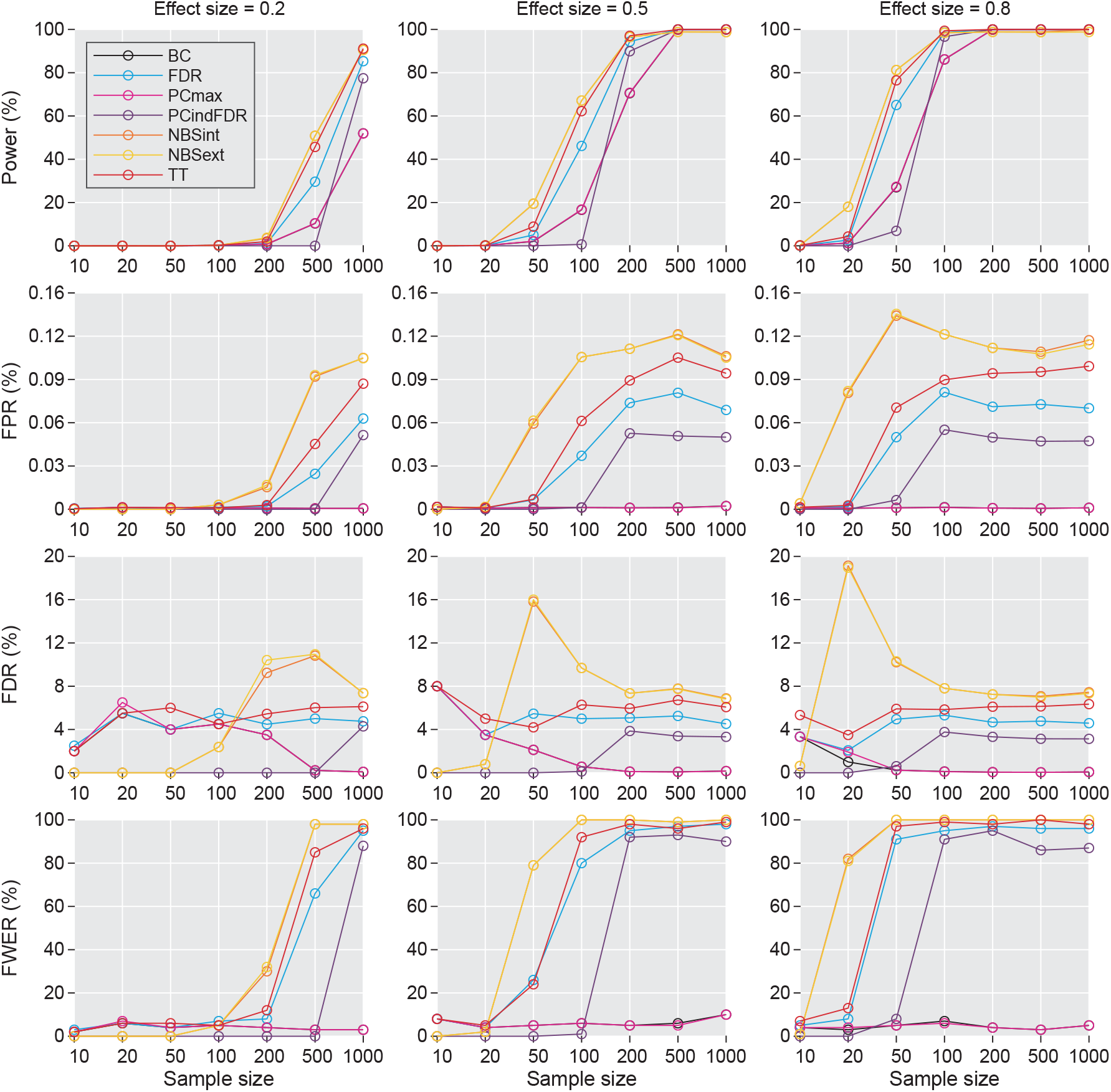
Benchmarking the TFNOS-TT procedure and six baselines. The TT procedure is compared to six baseline procedures for statistical performance (i.e., power, FPR, FDR, and strong-sense FWER) across three effect sizes and seven sample sizes (per group). The performance is almost identical between NBSint and NBSext, and between BC and PCmax, exhibiting nearly overlapping lines. See Section 2.5.3 for explanations of the code names of the procedures.

In fact, for a given effect size, if the sample size is too small, any procedure will be gravely underpowered, whereas a large enough sample size will provide adequate power (e.g., 80%) with all procedures. Intuitively, the cluster-level inferences were more powerful than the edge-level inferences. The weak-sense FWER-controlling procedures (e.g., NBSext, NBSint, TT, and FDR) exhibit greater power than strong-sense FWER-controlling procedures (e.g., PCmax and BC), except that PCindFDR may be limited by the resolution of the cluster-specific *p*-value. The NBSext and NBSint procedures exhibited the highest power at certain combinations of effect size and sample size but at the cost of discarding control of the FDR. For example, at the effect size of 0.8 and sample size of 20, the NBSint procedure has about 13.7% higher power than the TT procedure, but at the same time has 15.6% higher FDR. The power of the TT procedure was no less than the FDR for all combinations of effect sizes and sample sizes. It not only exhibits leading statistical power but can control FDR at a level (∼ 6%, see Fig. S11 for 95% confident intervals of the FDR) approaching that of the FDR procedure (nominal 5%). Given a total sample size, unbalanced sample sizes between groups will reduce power compared to the case of balanced sample sizes between groups (Fig. S12). In addition, the patterns of change in FDR and FWER with sample size or effect size for the seven statistical procedures can be categorized into two groups. The first group, comprising procedures with weak control of FWER (i.e., NBSint, NBSint, TT, FDR, and PCindFDR) tends to increase with effect size or sample size, whereas the other group, comprising procedures with strong control of FWER (i.e., BC and PCmax) maintain a remarkably low level across effect size and sample size. More formally, the 95% confidence intervals 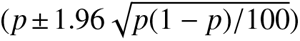 for the strong-sense FWER estimated by the BC and PCmax procedures at all combinations of effect sizes and sample sizes cover or slightly less than the nominal level of 5%.

Overall, our empirical results show that appropriate *E*/*H* parameters allow the TT procedure to maintain high statistical power while controlling the FDR at a low level. The TT procedure is superior to the NBS and conventional FDR procedures. Compared with the strong-sense FWER-controlling procedures, the TT procedure can markedly reduce the requirement for sample size^10^.

### 3.2. Experiment II: TFNOS-PC simulations

#### 3.2.1. Benchmarking cluster-level inference procedures

We included the six variants of the TFNOS-PC procedures (i.e., PCmax, PCmaxV, PCindBC, PCindBCV, PCindFDR, and PCindFDRV) in a benchmark with two other procedures (i.e., BC, FDR) for statistical performance (Section 2.6). The performance metrics for the eight procedures were calculated at the same spatial scale as the level of inference—at the cluster level. The eight procedures can be also divided into two groups:(1) FDR-controlling procedures (FDR, PCindFDC, and PCindF-DRV; controlling the FWER in the weak sense); (2) FWER-controlling procedures (BC, PCmax, PCmaxV, PCindBC, and PCindBCV; controlling the FWER in the strong sense).

As illustrated in Fig. 7, the three FDR-controlling procedures offered higher power than the five FWER-controlling procedures before reaching a power of 100%. In general terms, the FDR-controlling procedures exhibited higher FPR, FDR, and FWER in comparison to the FWER-controlling procedures. The estimated FDR of all eight procedures was smaller than 5%. Controlling the FWER in the strong sense will automatically control the FDR. As anticipated, the FDR-controlling procedures cannot control the FWER in a strong sense (Fig. 7). Of note, for procedures with cluster-specific null distributions, pooling edge-wise test statistics or signals within the predefined cluster yielded similar performance (PCindFDR vs. PCindFDRV and PCindBC vs. PCindBCV). PCindFDRV and PCindBCV dramatically reduced the number of tests and were much simpler compared to PCindFDR and PCindBC. Furthermore, in terms of the power of the nonparametric FWER-controlling procedures, PCmaxV showed almost the same performance compared to PCindBC and PCindBCV, but no additional corrections were required (using an overall null distribution); all of which outperformed PCmax. In this experimental setup, the assumptions of the edge-wise parameter tests are satisfied, and the BC (vs. PCindBC, PCindBCV, or PCmaxV) and FDR (vs. PCindFDR or PCindFDRV) are simpler procedures without compromise in performance.

**Fig. 7.**
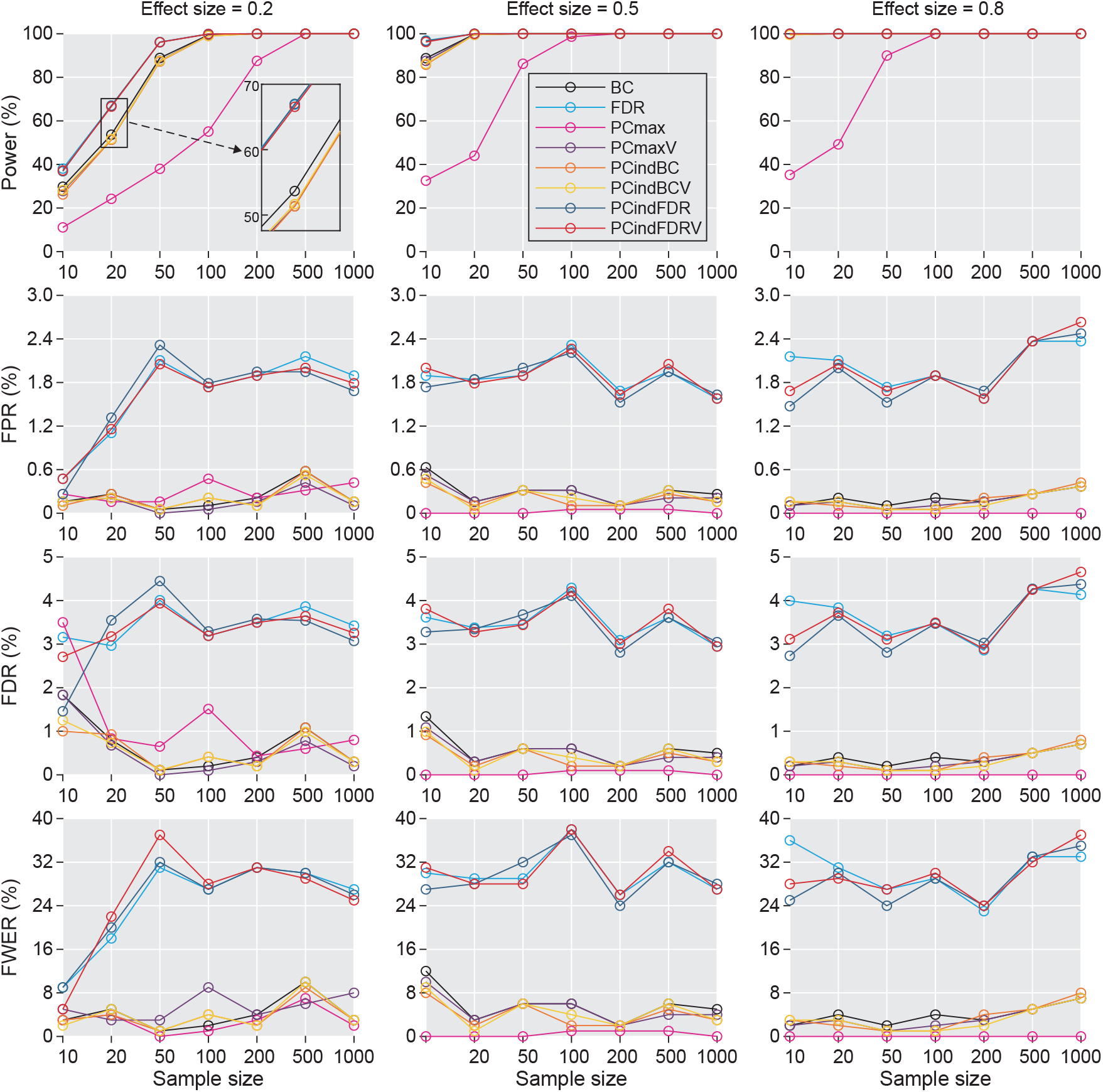
Benchmarking TFNOS-PC procedures and baselines. The performance of the TFNOS-PC procedures (six variants, i.e., PCmax, PCmaxV, PCindBC, PCindBCV, PCindFDR, and PCindFDRV) and two baseline procedures (i.e., BC and FDR) were evaluated across three effect sizes and seven sample sizes (per group).

### 3.3. Experiment III: Test in the fMRI dataset

Finally, we investigated the group-level difference in P-FC and CCM-EC between patients with SCI and HC, as illustrative examples of our TFNOS framework. Based on previous research (Ge et al., 2021), we considered that SCI alterations to brain networks may involve a large number of edges that may satisfy the implicit assumptions of both the TFNOS-TT and TFNOS-PC procedures. Given the sample size of the fMRI dataset, it implies that only the stronger effect can be detected with adequate power (Section 3.1.3). The PCindFDRV was chosen as a representation for the TFNOS-PC procedures.

The results of P-FC and CCM-CE from both the TT and PCindFDRV procedures confirmed that SCI patients exhibited large-scale “disrupted” interactions between brain regions compared to HC (Fig. S13 and Fig. 8). Additionally, it was observed that the number of edges with *p* < 0.05 resulting from the TT procedure was sensitive to the *E* and *H* parameters, underscoring the importance of selecting appropriate parameters. Here, we used *E* = 0.4 and *H* = 3.0, but it was noted that using *E* = 0.4 and *H* = 7.0 would yield more conservative results (Figs. 4 and 8). Furthermore, SCI-induced significant alterations (*p* < 0.05) in the strength of causative effect were with a considerable portion of bidirectional (57% in TT; 99% in PCindFDRV) (Fig. 8). The PCindFDRV has less spatial specificity. See results for another TFNOS-PC variant (PCmaxV) in Fig. S14. Roughly, our findings appeared to indicate that edges associated with the temporal lobe, limbic system, and temporal lobe exhibited larger observed effects.

**Fig. 8.**
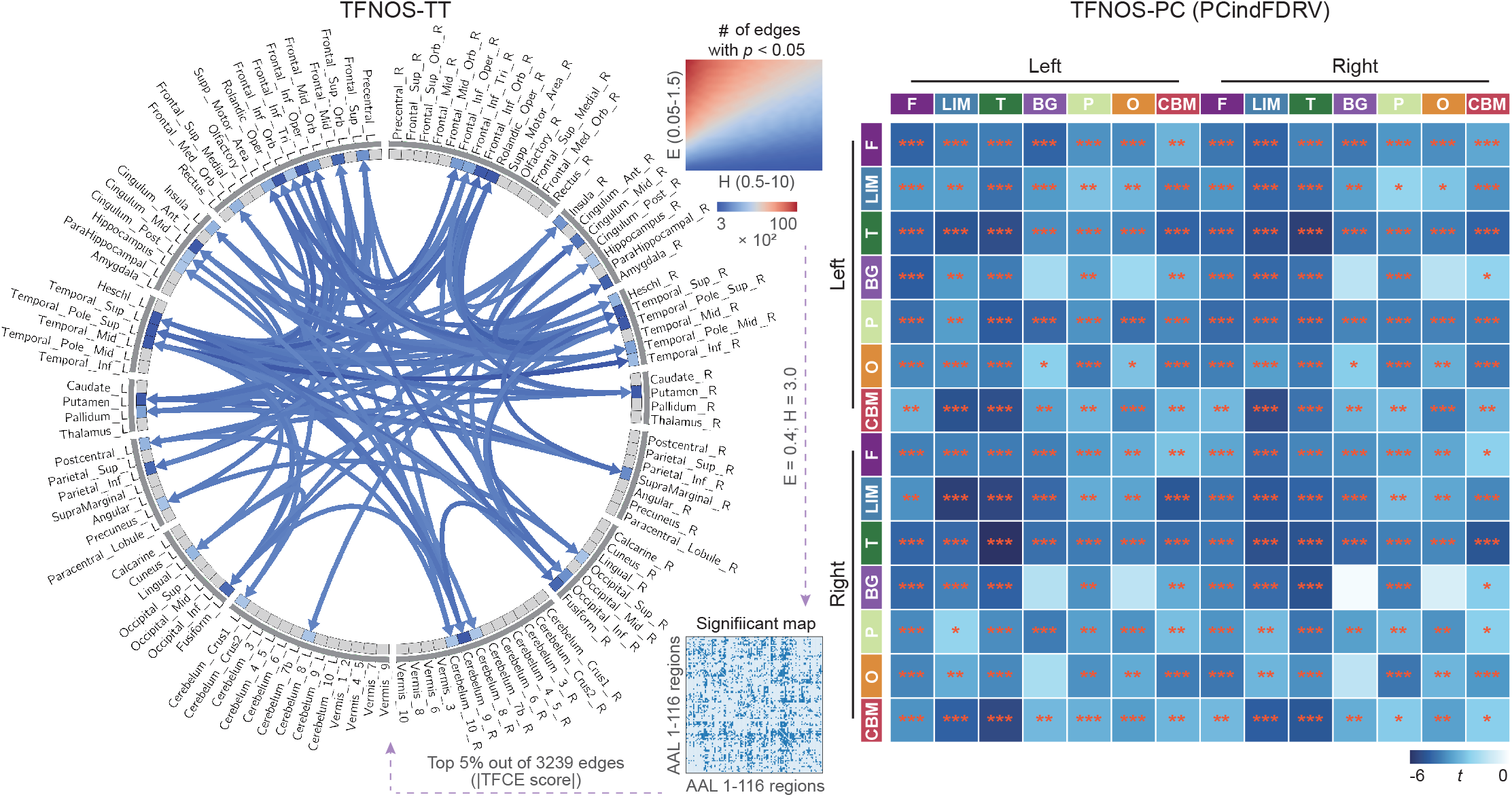
Using the TFNOS framework to identify differences in CCM-based directed brain networks between patients with SCI and HC. (Left) Using TFNOS-TT procedure with *E* = 0.4 and *H* = 3.0. For visualization purposes, the Circos plot only shows edges with *p* < 0.05 where the absolute value of the TFCE score is in the top 5% (out of the 3239 edges with *p* < 0.05). The blue lines indicate SCI < HC, with darker colors representing larger absolute TFCE scores. (Right) Using TFNOS-PC procedure (PCindFDRV). \**p* < 0.05; \*\**p* < 0.01; \*\*\**p* < 0.001. F = frontal; LIM = limbic; T = temporal; BG = basal ganglia; P = parietal; O = occipital; CBM = cerebellum.

## 4. Discussion

Collectively, we employed the proposed TFNOS framework to systematically evaluate the performance of the threshold-free statistical procedures across different spatial topologies of effects, effect sizes, and sample sizes. The TFNOS framework reviews and expands current relevant nonparametric procedures, enabling their broad application in various contexts on group-level statistical inferences in brain networks. Our empirical findings demonstrate that the TFNOS is a powerful framework with many degrees of freedom, and provide empirical and principled guidance. While our subsequent discussions primarily focus on inferences related to brain network edges, the key points are also applicable to statistical inferences concerning brain network nodes.

The TFNOS-TT procedure is characterized as being threshold-free, but not parameter-free (Baggio et al., 2018; Vinokur et al., 2023). Our research emphasizes the importance of selecting appropriate *E* and *H* parameters to achieve leading power while maintaining low type I error rates. Based on our study, we recommend utilizing *E* values of 0.4 and *H* values ranging from 3.0 (more liberal) to 7.0 (more conservative). (Fig. 4). Our empirical evidence demonstrates that setting *E* and *H* to these recommended values can effectively control the FDR (e.g., *H* = 3.0, FDR < 10%; *H* = 7.0, FDR < 5%). This is validated across different effect topologies, effect sizes, and sample sizes (Figs. 4–6, Fig. S11, and Fig. S12).

Conversely, our findings suggest that the recommended values of *E* = 0.5 and *H* = 2.0, derived from MRI voxels (Smith and Nichols, 2009), should not be simply generalized to typical brain network contexts, as this may lead to a high number of false positives (e.g., FDR > 20%) (Fig. 4). Moreover, the previously suggested values of *E* = 0.5 (*H* = 2.25–3.0) or *E* = 0.75 (*H* = 3.0–3.5) from previous TFNBS research (Baggio et al., 2018) were found to be overly liberal in our experimental setup. For example, the FDR would even be greater than 50% when *E* = 0.75 (*H* = 3.0–3.5) (Fig. 4). We attribute these discrepancies to differences in test statistics used (see Section 2.5.1) and the focus on statistical power and FPR in previous research (Baggio et al., 2018), without considering the FDR. Note that the recommended *E* and *H* values for brain network edges may not be universally applicable to all brain network-related research contexts. For instance, in statistical inference on EEG sensor-level measures, the recommended values are different, with *E* = 1 and *H* = 2.0–3.0 being more appropriate (Figs. S8-S10), which is similar to previous simulation results on EEG source space dipoles (Mensen and Khatami, 2013) and satisfies the TFCE default parameter settings of the PALM tool^11^ for surface data. Some studies have used the NBS procedure with different cluster-forming thresholds to show that the existence of an effect is not deliberately screened for (Ge et al., 2022, 2021). We provide here a suitable alternative procedure, while substantially alleviating previous concerns that the TFNBS is not “parameter-free” (Vinokur et al., 2023). The TFNOS-TT procedure may also have advantages in terms of test-retest reliability (Chen et al., 2018). In general, our findings offer empirical evidence and guidelines for setting *E* and *H* parameters in open toolboxes (Nieto-Castanon, 2020; Tournier et al., 2019; Gramfort et al., 2013; Oostenveld et al., 2011) commonly used and future related research.

In numerical simulations, the precise locations of the ground-truth effects are known, providing the opportunity to estimate performance metrics at the edge level for the TFNOS-TT and NBS procedures (see Section 2.4). However, this necessitates a strong “assumption”: for TFNOS-TT inference, an edge with a *p*-value less than 0.05 is considered significant, while for NBS inference, all edges within a significant cluster are considered significant. Such empirical evaluation of performance is not new and follows a similar approach in previous work on the NBS, TFNBS, and others (Zalesky et al., 2010; Baggio et al., 2018; Noble et al., 2020, 2022; Zhang et al., 2022). When cluster-level TFNOS-TT and NBS inferences were interpreted at the edge level, their statistical power surpassed that of edge-level inferences (Fig. 6). Intriguingly, our empirical results show that the TFNOS-TT procedure can maintain stable control of the FDR at a specific level by using the recommended values of the *E*/*H* parameter (Figs. 4-6), whereas the NBS cannot (Fig. 6). For example, the FDR was all ≤ 5% as depicted in Fig. 5 and gradually stabilized at around 6% with increasing sample size as shown in Fig. 6 and Fig. S11. We speculate that this “property” of TFNOS-TT is partly attributed to its preservation of more spatial information than traditional clustering inference, e.g., the “local maxima” of the output TFCE score map are at the same locations as those in the original statistic map (Smith and Nichols, 2009; Spisák et al., 2019). The ability of the TFNOS-TT procedure to control FDR is appealing, although not theoretically guaranteed, and may enhance the spatial specificity (localization power) of the TFNOS-TT inference. This topic deserves deep investigation in the future.

The TFNOS framework offers a generalized framework for group-level threshold-free statistical inference in the field of network neuroscience. The selection of a specific statistical procedure for a given study is contingent upon various factors, including the spatial structure of effects, effect size, sample size, test statistic of interest, and research objectives. While generic procedures like BC and FDR can yield sufficient statistical power to identify significant effects with a large enough sample size (Figs. 6 and 7), brain network studies often have relatively small sample sizes (Helwegen et al., 2023). Moreover, the true effect sizes are unknown, requiring the estimation and determination of the minimal effect sizes of interest (Lakens, 2022; Helwegen et al., 2023). The TFNOS-TT and TFNOS-PC procedures implicitly make assumptions about the spatial structure of the effect of interest. The TFNOS-TT procedure assumes that the edges with the effect of interest can form specific spatial extended components (e.g., WCC, SCC, and clique), while the TFNOS-PC procedure assumes that the effect of interest is broadly distributed in (specific) predefined clusters with consistent signs/directions of effects within a cluster. These assumptions are not mutually exclusive and may both be satisfied in certain situations. Generally, the TFNOS-TT procedures are suitable for scenarios with effect topologies resembling star, mesh, tree, or their compositions (Ge et al., 2022, 2021; Chen et al., 2023), while the TFNOS-PC procedures are suitable for scenarios with widespread spatial effects, such as large-scale alterations of the brain network induced by severe brain injury or tasks (Sitt et al., 2014; Hao et al., 2023a; Noble et al., 2022). The TFNOS framework may be less sensitive to focal effects, particularly for the TFNOS-PC procedures, where large focal effects might only represent small cluster-level effects when averaged across all edges in a predefined cluster. Our empirical findings suggest that the TFNOS-TT can control the FDR at the edge level, to some extent alleviating concerns about the spatial specificity paradox (Rosenblatt et al., 2018; Goeman et al., 2023). Note that controlling FDR at γ level (e.g., 5%) does not guarantee that the false discovery proportion must be less than or equal to γ with a high probability (Dmitrienko et al., 2009). The TFNOS-TT procedures and the FDR-controlling procedures (PCindFDR and PCindFDRV) in TFNOS-PC are more suitable for exploratory rather than confirmatory studies.

The TFNOS is a nonparametric statistical framework based on the permutation test and offers many degrees of freedom. The TFNOS-TT procedure provides flexibility in defining clusters, allowing for consideration of factors such as the topology of the effect of interest and knowledge experience (Fig. 1). The TFNOS-PC procedures encompass several variants, some of which are very simple (e.g., PCmaxV, PCindBCV, PCindFDRV), which allows one to choose any test statistic one considers appropriate. Additionally, The determination of predefined clusters is based on predefined node groups, and it is crucial to establish the division of node groups on physiological grounds, as arbitrary divisions can significantly impact statistical performance, for instance, by treating each node as a group (Fig. 6). Moreover, the PCmaxV procedure utilizes the maximum statistics of all clusters to construct the empirical null distribution to strongly control FWER but at the cost of reducing the power of clusters with smaller statistics (Maris and Oostenveld, 2007). In contrast, the PCindBCV procedure constructs cluster-specific null distributions but requires multiple testing corrections to control the false positives (Noble and Scheinost, 2020). The comparative advantage in terms of statistical power between both approaches is not theoretically clear. However, the PCindBCV requires a larger number of permutations to improve the *p*-value resolution, when the number of clusters is relatively large. The PCmaxV involves averaging the edge values within a cluster, which reduces variance and consequently increases the cluster-level effect size, potentially explaining the power difference between PCmaxV and PCmax. Furthermore, this study presents an illustrative example of applying the framework to real data (Fig. 8). While our results were in line with the previous findings using the NBS (Ge et al., 2021), we got rid of the dependence on the cluster-forming threshold and provided better error rate control beyond only weak control of FWER (Zalesky et al., 2010). For TFNOS-PC procedures using cluster-specific null distributions, control FWER in the strong sense can also be achieved using procedures based on the closure principle other than traditional Bonferroni correction (Marcus et al., 1976; Dmitrienko et al., 2009). The TFNOS framework can be easily generalized to a variety of other experimental designs, modalities, and spatial scales and offers a comprehensive set of statistical methods that enhance the capacity to capture diverse effects in brain networks. It may also have the potential to be applied or extended to other biomedical and network science fields that face similar issues to brain networks.

The current study has certain limitations and potential for future expansion. We used simulated data sampled from a normal distribution, and the distribution type may be presented differently in the various contexts of real data. Moreover, while the impact of spatial dependence on statistical performance was assessed, it is essential to validate these findings with higher-dimensional FC. The recently released connectomes for 40000 UK Biobank participants offer an opportunity for further investigation in this regard (Mansour L. et al., 2023). In numerical simulations, we assigned a fixed effect size *d* (observed effects are variable) to all edges set as ground truth. Future work could consider setting mixed effect sizes for ground truths, although the distribution of effect sizes remains uncertain across scenarios. Furthermore, our simulation-based results illustrate a trend in the power of different statistical procedures to vary with sample size or effect size. However, these findings should not be directly extrapolated to estimate sample sizes in various scenarios, especially when they distinctly diverge from our experimental settings.

## 5. Conclusions

The growing interest in finer-grained network explorations has been accompanied by increasing concerns about the reproducibility of the results and statistical power. Here, building upon the work of others, we introduce a threshold-free brain network cluster-level statistical inference framework, TFNOS, which encapsulates statistical pipelines that are free from the selection of a hard cluster-formation threshold as in the traditional NBS procedure. We demonstrate the performance of the framework under different effect topologies, sample sizes, and effect sizes through numerical simulations. Results indicate that the TFNOS framework performs excellently across a wide range of statistical procedures. The framework can be used in various scenarios regarding group-level statistical inference on nodes or edges of the brain network. The present work tends to provide empirical and principled criteria to the population regarding the choice of statistical procedures, thereby contributing to increased reproducibility and sensitivity in future research.

## 6. Data and code availability statements

The fMRI and EEG data cannot be shared due to ethics committee restrictions. The main codes used to generate simulations and figures are publicly available at https://osf.io/vxmd4/.

### CRediT authorship contribution statement

**Zexuan Hao:** Conceptualization, Data curation, Formal analysis, Investigation, Methodology, Software, Visualization, Writing – original draft, Writing – review & editing. **Pei Wang:** Validation, Writing – review & editing. **Xiaoyu Xia:** Data curation, Investigation, Resources, Writing – review & editing. **Yu Pan:** Data curation, Funding acquisition, Resources, Writing – review & editing. **Weibei Dou:** Conceptualization, Funding acquisition, Project administration, Resources, Supervision, Validation, Writing – review & editing.

### Declaration of competing interest

The authors declare no competing interests.

## Acknowledgements

This work was supported in part by the National Key Research and Development Program of China (2022YFC3601100 and 2022YFC3601105). We are grateful to Dr. Yunxiang Ge for pre-processing the fMRI data, and Dr. Yang Bai, Dr. Yong Wang, Peng Bo, and Yanlin Zhang for acquiring the EEG data. We thank the editors and anonymous reviewers in advance for their comments and suggestions.

## Footnotes

1 For instance, nodes can represent voxels, neuronal populations, brain regions, or sensors; edges can represent adjacency/neighbor relationships, neural fiber bundle connections, or functional interactions between nodes. Note that we do not distinguish between the terms {network, node, link} in network science and {graph, vertex, edge} in graph theory. We prefer to use the terms {network, node, edge}.

2 The FPR is defined as the proportion of true null hypotheses that are falsely rejected. The FWER is defined as the probability of at least one rejection of a true null hypothesis. The FDR is defined as the expected value of false discovery proportion (FDP; the proportion of rejected hypotheses that are falsely rejected). The FDP is sometimes referred to as observed FDR (Nichols and Hayasaka, 2003). In addition, the power can be referred to as sensitivity and 1− false negative rate (or type II error rate). The FPR can be referred to as Type I error rate and 1 − specificity.

3 Weak control of FWER just requires that the probability of rejecting one or more true null hypotheses is controlled less than the test level α under the complete null (“omnibus”) hypothesis (i.e., all null hypothesis are true); strong control of FWER requires that the probability of falsely rejecting at least one true null hypotheses is controlled less than α over any subset of the true null hypotheses, independently of whether the null hypothesis holds elsewhere.

4 The term “cluster” refers to a set of edges/nodes, implying that edges/nodes within the same cluster are more similar to each other in some aspects than to those in other clusters. Note that individual nodes/edges and the whole brain are extreme special cases of clusters. Unless otherwise noted, the cluster is the regular form.

5 An open MATLAB package of the TFCE developed by Dr. Mark Thornton is available at https://github.com/markallenthornton/MatlabTFCE. He also provided meaningful remark on the TFCE procedure at http://markallenthornton.com/blog/matlab-tfce/.

6 For a given test, the power is as a function of significant level α, effect size, and sample size. The reader can refer further to pp. 14–16 of Cohen (1988) if needed. Dr. Kristoffer Magnusson shows a nice interactive example at https://rpsychologist.com/d3/nhst/.

7 The number of possible permutations for different designs can be found at https://fsl.fmrib.ox.ac.uk/fsl/fslwiki/Randomise/Theory.

8 A derivation of Cohen’s *d* for the two independent samples case (Cohen, 1988) is as follows. *d* = *m*_*A*_ *m*_*B*_ /*σ*, where *m*_*A*_ and *m*_*B*_ are the mean of population A and B, respectively; σ is the standard deviation of either population, since σ_*A*_ and *σ*_*B*_ are assumed equal (i.e., *σ* = *σ*_*A*_ = *σ*_*B*_), and the pooled stan-dard deviation is 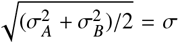 The effect size observed in the samples is an estimate of the real effect size.

9 The assumption behind the requirement that the covariance matrix is positive definite is that each edge contributes more or less independent information to some degree.

10 Under this experimental setup, we estimated the sample sizes (per group) required to achieve adequate power (80%) for the TT and BC procedures at different effect sizes. Specifically, the minimum sample size required approximately 800 vs. 1400 (TT vs. BC) subjects for *d* = 0.2, 130 vs. 230 subjects for *d* = 0.5, and 55 vs. 90 subjects for *d* = 0.8.

11 https://fsl.fmrib.ox.ac.uk/fsl/fslwiki/PALM.

## Notes

### Competing Interest Statement

The authors have declared no competing interest.

### Summary of Updates

The text was carefully revised; added two figures (Fig. 2 and 3) of pseudocodes on algorithms; original Figs. 1 and 6 revised; supplemental files updated.

