## Supplementary Material for "Threshold-Free Network-Oriented Statistics in Neuroscience"

### Supplementary Material for
### "Threshold-Free Network-Oriented Statistics in Neuroscience"

Hao *et al.*

**Algorithm:** TFNOS-TT for nodes

**Input :** Extension enhancement parameter $E$, height enhancement parameter $H$, # of thresholding steps $n$, # of permutation tests $T$, edge-wise statistical test $f$, cluster definition method $g$, nodal relationship matrix $\mathbf{A} \in \mathbb{R}^{C \times C}$, features across $C$ nodes of $N$ subjects $\mathbf{Y} \in \mathbb{R}^{N \times C}$, and design matrix $\mathbf{X} \in \mathbb{R}^{N \times M}$ with $M$ variables of interest (can only contain group indicator variable).

**Output:** Corrected $p$-values: $\boldsymbol{p} \in \mathbb{R}^C$, observed statistics $\boldsymbol{s}_o \in \mathbb{R}^C$, and TFCE scores $\boldsymbol{q}_o \in \mathbb{R}^C$.

/* Observed statistics and TFCE scores */

$\boldsymbol{s}_o = f(\mathbf{Y}, \mathbf{X})$;

$\boldsymbol{q}_{o,p} \leftarrow$ `get_TFCE_score`$(\boldsymbol{s}_o, \mathbf{A}, E, H, n, g)$;

$\boldsymbol{q}_{o,n} \leftarrow$ `get_TFCE_score`$(-\boldsymbol{s}_o, \mathbf{A}, E, H, n, g)$;

$\boldsymbol{q}_o \leftarrow \mathrm{abs}(\boldsymbol{q}_{o,p} + \boldsymbol{q}_{o,n})$;

/* Nonparametric permutation testing */

$t \leftarrow 1$;

$\boldsymbol{n\_count} \leftarrow \mathbf{1} \in \mathbb{R}^C$;

**while** $t \leq T$ **do**

    $t \leftarrow t + 1$;

    $\mathbf{X}_r \leftarrow$ `random_shuffle`$(\mathbf{X})$; // e.g., orders, labels

    $\boldsymbol{s}_r = f(\mathbf{Y}, \mathbf{X}_r)$;

    $\boldsymbol{q}_{r,p} \leftarrow$ `get_TFCE_score`$(\boldsymbol{s}_r, \mathbf{A}, E, H, n)$;

    $\boldsymbol{q}_{r,n} \leftarrow$ `get_TFCE_score`$(-\boldsymbol{s}_r, \mathbf{A}, E, H, n)$;

    $\boldsymbol{q}_r \leftarrow \mathrm{abs}(\boldsymbol{q}_{r,p} + \boldsymbol{q}_{r,n})$;

    $\boldsymbol{n\_count} \leftarrow \boldsymbol{n\_count} + \max(\boldsymbol{q}_r) \geq \boldsymbol{q}_o$;

/* Corrected $p$-values */

$\boldsymbol{p} \leftarrow \boldsymbol{n\_count}/(T+1)$;

**Function** `get_TFCE_score`$(\boldsymbol{s}, \mathbf{A}, E, H, n, g)$**:**

    $\boldsymbol{q} \leftarrow \mathbf{0} \in \mathbb{R}^C$ ;

    $dh \leftarrow \max(\boldsymbol{s})/n$;

    **if** $dh > 0$ **then**

        **for** $h \leftarrow dh : dh : \max(\boldsymbol{s})$ **do**

            $\boldsymbol{mask} \leftarrow \boldsymbol{s} < h$;

            $\mathbf{A}(\boldsymbol{mask}, \boldsymbol{mask}) \leftarrow 0$;

            $cluster\_num, \boldsymbol{node\_set}, \boldsymbol{size} \leftarrow$ `get_cluster`$(\mathbf{A}, g)$;

            **for** $c \leftarrow 1 : cluster\_num$ **do**

                $\boldsymbol{q}(\boldsymbol{node\_set}(c)) \leftarrow \boldsymbol{q}(\boldsymbol{node\_set}(c)) + \boldsymbol{size}(c)^E \times h^H \times dh$;

    **return** $\boldsymbol{q}$;

**Fig. S1.** Pseudocodes of the TFNOS-TT procedure for network nodes.

**Algorithm:** TFNOS-PC for nodes

**Input :** # of permutation tests $T$, statistical test $f$, cluster definition method $g$, features across $C$ nodes of $N$ subjects $\mathbf{Y} \in \mathbb{R}^{N \times C}$, and design matrix $\mathbf{X} \in \mathbb{R}^{N \times M}$ with $M$ variables of interest (can only contain group indicator variable).

**Output:** Corrected $p$-values: $\boldsymbol{p} \in \mathbb{R}^K$ and observed statistics $\boldsymbol{s}_o \in \mathbb{R}^K$ for $K$ predefined node groups, $K$ depends on the $g$.

/* Observed cluster-wise statistics */

$\boldsymbol{s}_o \leftarrow$ `get_PC_stats`$(\mathbf{Y}, \mathbf{X}, f, g)$;

$\boldsymbol{s}_o \leftarrow \text{abs}(\boldsymbol{s}_o)$;

/* Nonparametric permutation testing */

$t \leftarrow 1$;

$\boldsymbol{n\_count} \leftarrow \mathbf{1} \in \mathbb{R}^M$;

**while** $t \leq T$ **do**

- $t \leftarrow t + 1$;
- $\mathbf{X}_r \leftarrow$ `random_shuffle`$(\mathbf{X})$; // e.g., orders, labels
- $\boldsymbol{s}_r \leftarrow$ `get_PC_stats`$(\mathbf{Y}, \mathbf{X}_r, f, g)$;
- $\boldsymbol{s}_r \leftarrow \text{abs}(\boldsymbol{s}_r)$;
- /* Option 1: an overall null distribution */
- $\boldsymbol{n\_count} \leftarrow \boldsymbol{n\_count} + \max(\boldsymbol{s}_r) \geq \boldsymbol{s}_o$;
- /* Option 2: cluster-specific null distribution */
- $\boldsymbol{n\_count} \leftarrow \boldsymbol{n\_count} + \boldsymbol{s}_r \geq \boldsymbol{s}_o$

/* Corrected $p$-values */

$\boldsymbol{p} \leftarrow \boldsymbol{n\_count}/(T + 1)$;

$\boldsymbol{p} \leftarrow$ `FWER_or_FDR_correction`$(\boldsymbol{p})$ ; // no need for option 1

**Function** `get_PC_stats`$(\mathbf{Y}, \mathbf{X}, f, g)$**:**

- /* Option 1: get node-wise test statistics first */
- $\boldsymbol{s}_{ori} = f(\mathbf{Y}, \mathbf{X})$;
- $\boldsymbol{s} \leftarrow$ `get_cluster_stats`$(\boldsymbol{s}_{ori}, g)$;
- /* Option 2: get cluster-wise signals first */
- $\mathbf{Y}_c, \mathbf{X}_c \leftarrow$ `get_cluster_signal`$(\mathbf{Y}, \mathbf{X}, g)$;
- $\boldsymbol{s} = f(\mathbf{Y}_c, \mathbf{X}_c)$;
- **return** $\boldsymbol{s}$;

**Fig. S2.** Pseudocodes of TFNOS-PC procedures for network nodes.

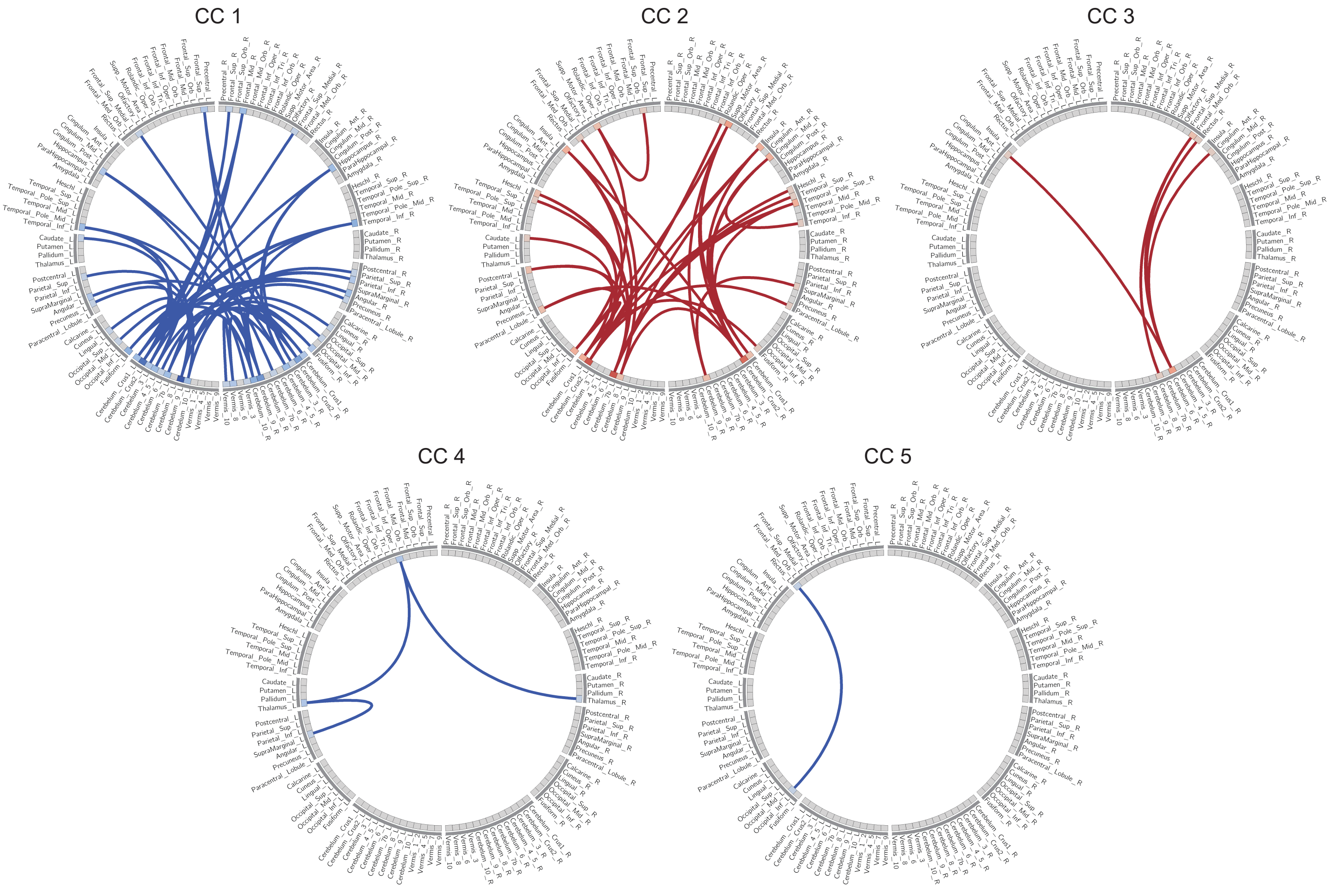


**Fig. S3.** Ground-truth connected components (CC) in Experiment I for parameter search. Schematic representation of the five CCs used as ground-truth effects. Edges in CC 1, CC4, and CC 5 were set with negative effects, while CC 2 and CC 3 were set with positive effects. See more details in the main text (Section 2.5.1).

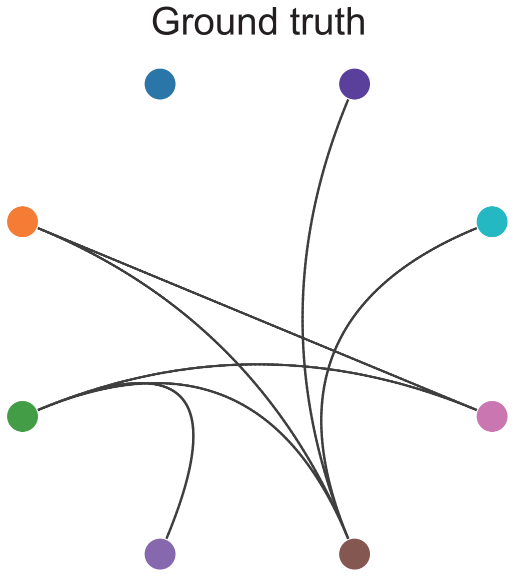


**Fig. S4.** Ground-truth setup in Experiment I for investigating the impact of spatial dependence. The shown edges were set with ground-truth effects. See more details in the main text (Section 2.5.2).

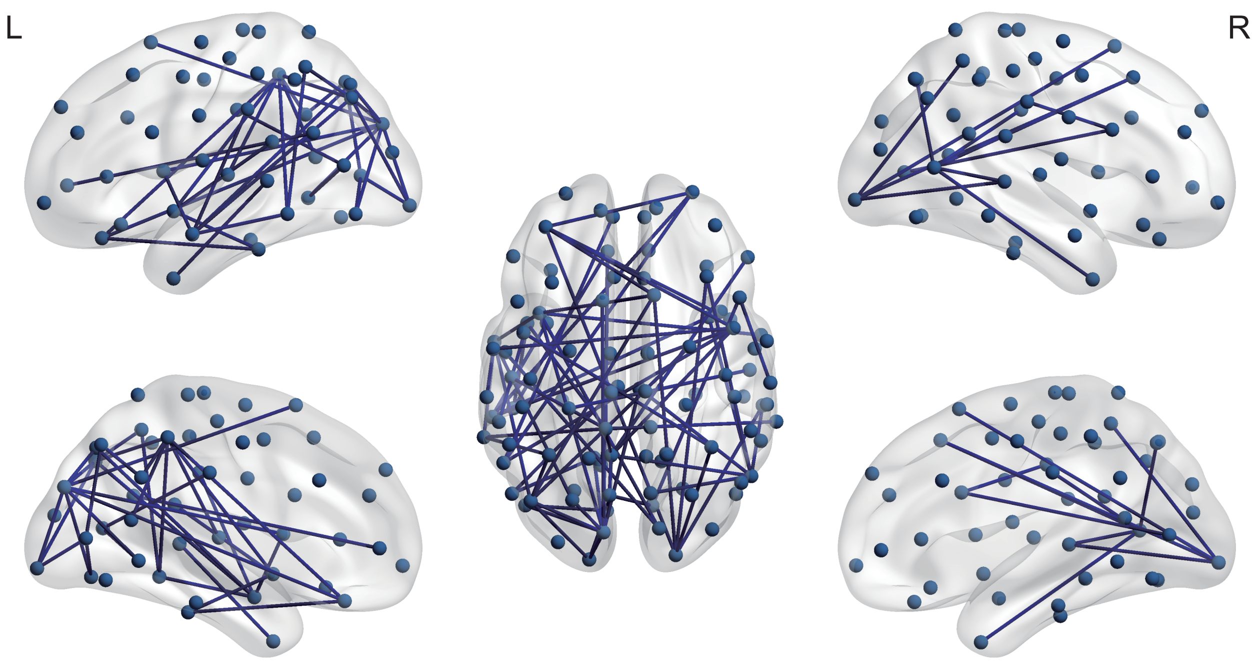


**Fig. S5.** Ground-truth setup in Experiment I for benchmarking performance of the TFNOS-TT procedure and corresponding baselines. See more details in the main text (Section 2.5.3). The graphs were created using the BrainNet Viewer toolbox (Xia et al., 2013).

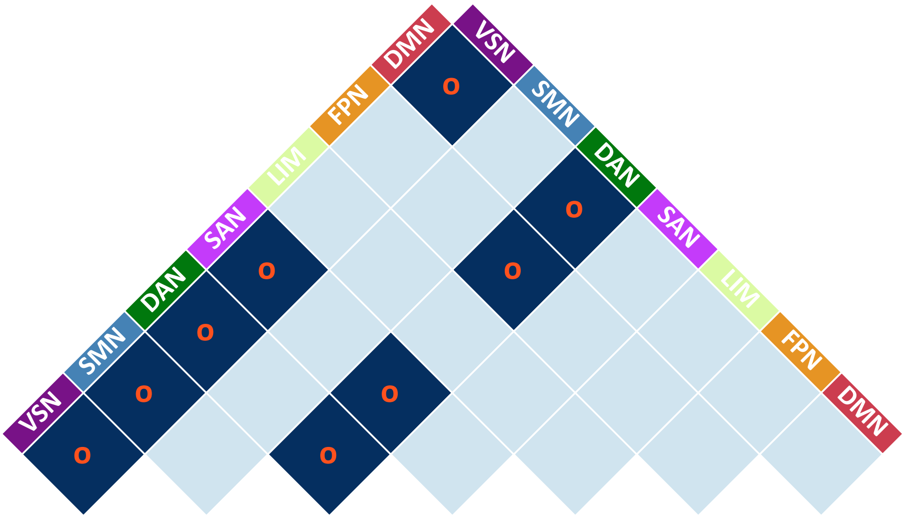


**Fig. S6.** Ground-truth setup in Experiment II for benchmarking cluster-level inference procedures. The nine subnetworks marked by "O" were set with ground-truth effects. VSN = visual network; SMN = sensorimotor network; DAN = dorsal attention network; san = salience attention network; LIM = limbic network; FPN = frontoparietal network; DMN = default mode network. See more details in the main text (Section 2.6).

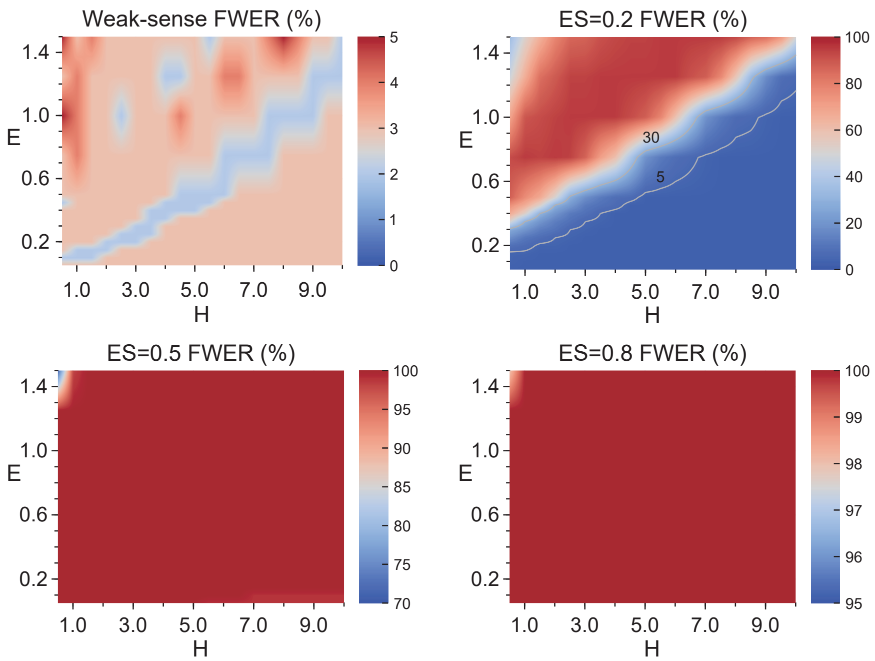


**Fig. S7.** FWER heatmaps. Heatmaps display empirical FWER of the TFNOS-TT procedure (edges) under numerical simulations (100 repetitions) for 600 $E/H$ parameter combinations (30 $E$ and 20 $H$ values), with effect size (Cohen's $d$) of 0, 0.2, 0.5, and 0.8. For better visualization, the data for the heatmaps were linearly interpolated somewhat and some necessary contours were added (in gray). The results show that the TFNOS-TT procedure controls the FWER in the weak sense. ES = effect size.

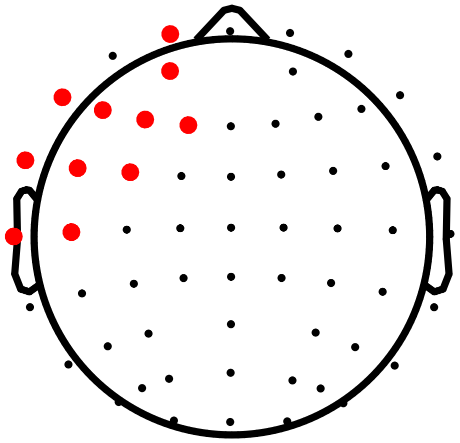

**Fig. S8.** Ground-truth setup in Experiment I for parameter search of the TFNOS-TT procedure for nodes. Here nodes represent EEG electrodes. We first extracted the relative power in the alpha band (8–13 Hz) from eyes-closed resting-state EEG data of 45 left-sided stroke patients and 26 healthy individuals (Hao et al., 2023). We then performed node-wise two-sample $t$-test and converted the $t$-values to observed effect sizes $d$ ($|t| \times \sqrt{1/n1 + 1/n2}$). Finally, nodes with suprathreshold ($d > 0.8$) observed effect sizes were set with ground-truth effects. See Hao et al. (2023) for data acquisition and preprocessing. See Fig. S1 of Hao et al. (2023) for the setting of neighboring relations between electrodes.

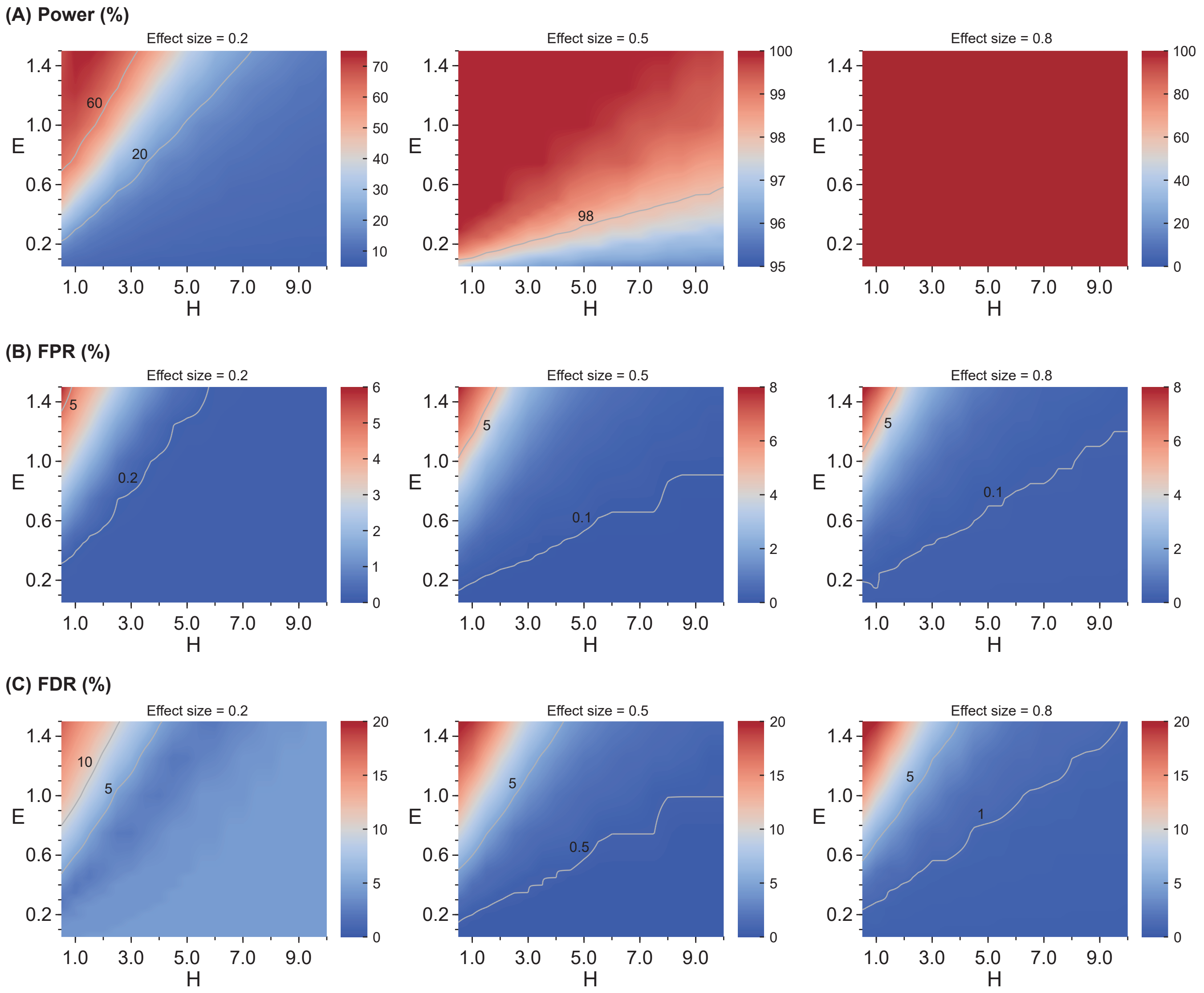


**Fig. S9.** Parameter search of TFNOS-TT procedure for nodes. Heatmaps display empirical power (top), FPR (middle), and FDR (bottom) under numerical simulations (100 repetitions) for 600 $E/H$ parameter combinations (30 $E$ and 20 $H$ values), with effect size (Cohen's $d$) of 0.2 (left), 0.5 (middle), and 0.8 (right). For better visualization, the data for the heatmaps were linearly interpolated somewhat, and some necessary contours were added (in gray). See Fig. S10 for the heatmaps of FWER.

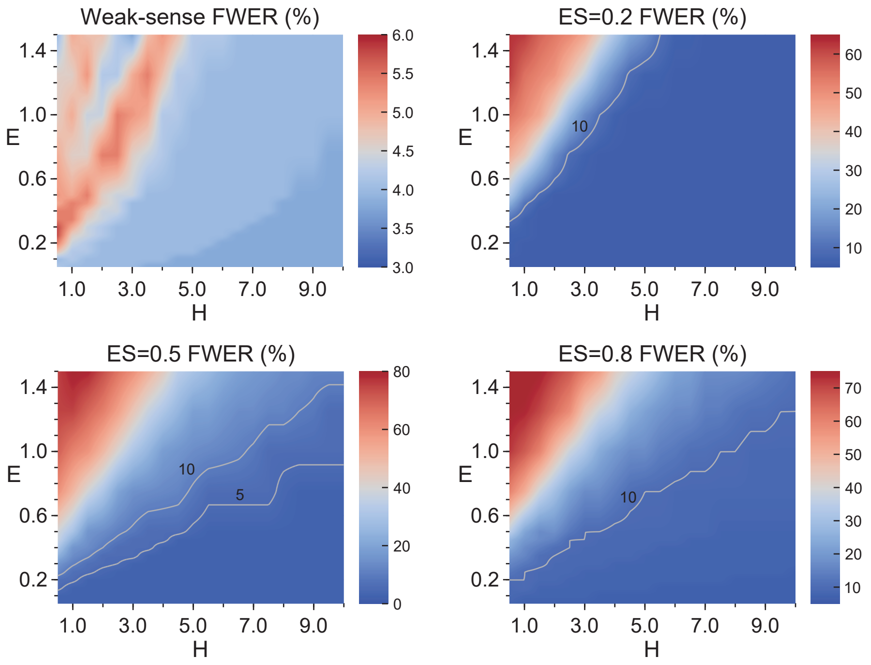


**Fig. S10.** FWER heatmaps. Heatmaps display empirical FWER of the TFNOS-TT procedure (nodes) under numerical simulations (100 repetitions) for 600 $E/H$ parameter combinations (30 $E$ and 20 $H$ values), with effect size (Cohen's d) of 0, 0.2, 0.5, and 0.8. For better visualization, the data for the heatmaps were linearly interpolated somewhat and some necessary contours were added (in gray).

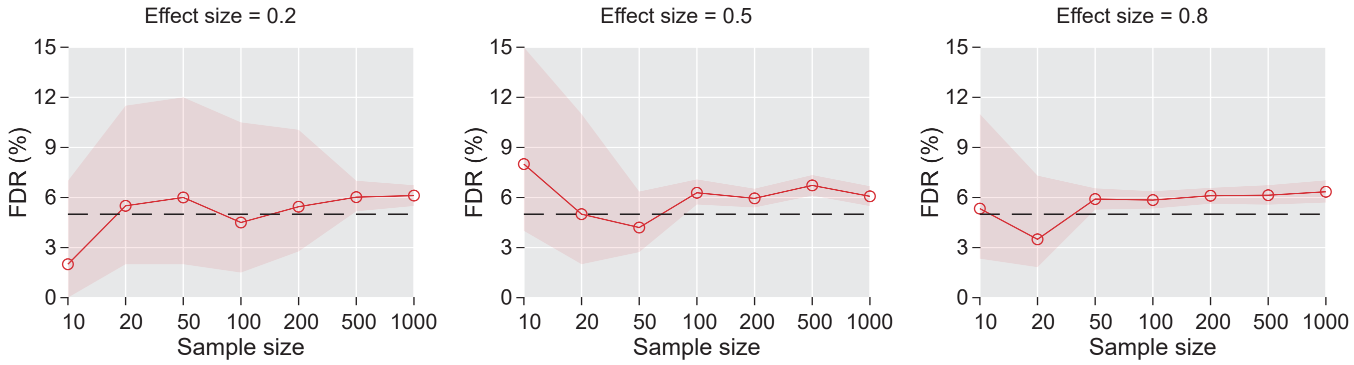


**Fig. S11.** The FDR and 95% confidence intervals of the TFNOS-TT procedure in Experiment II. The FDR (95% confidence intervals) was estimated across three effect sizes and seven sample sizes (per group). Bootstrap confidence intervals were computed using a bias-corrected and accelerated percentile method with 10000 resamples.

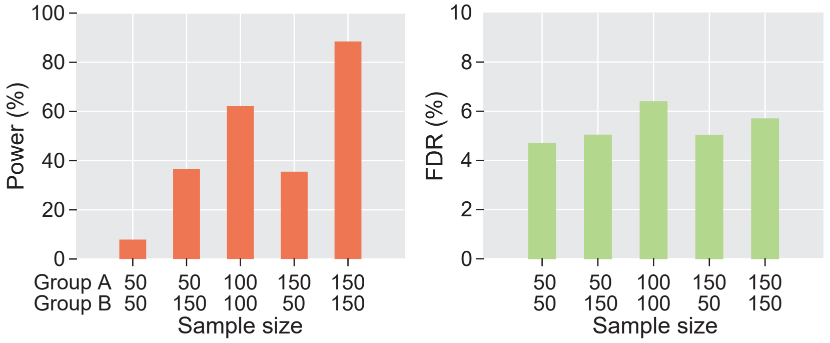


**Fig. S12.** Examples (TFNOS-TT, $d$ = 0.5) of power and FDR when sample sizes are the same or different between groups.

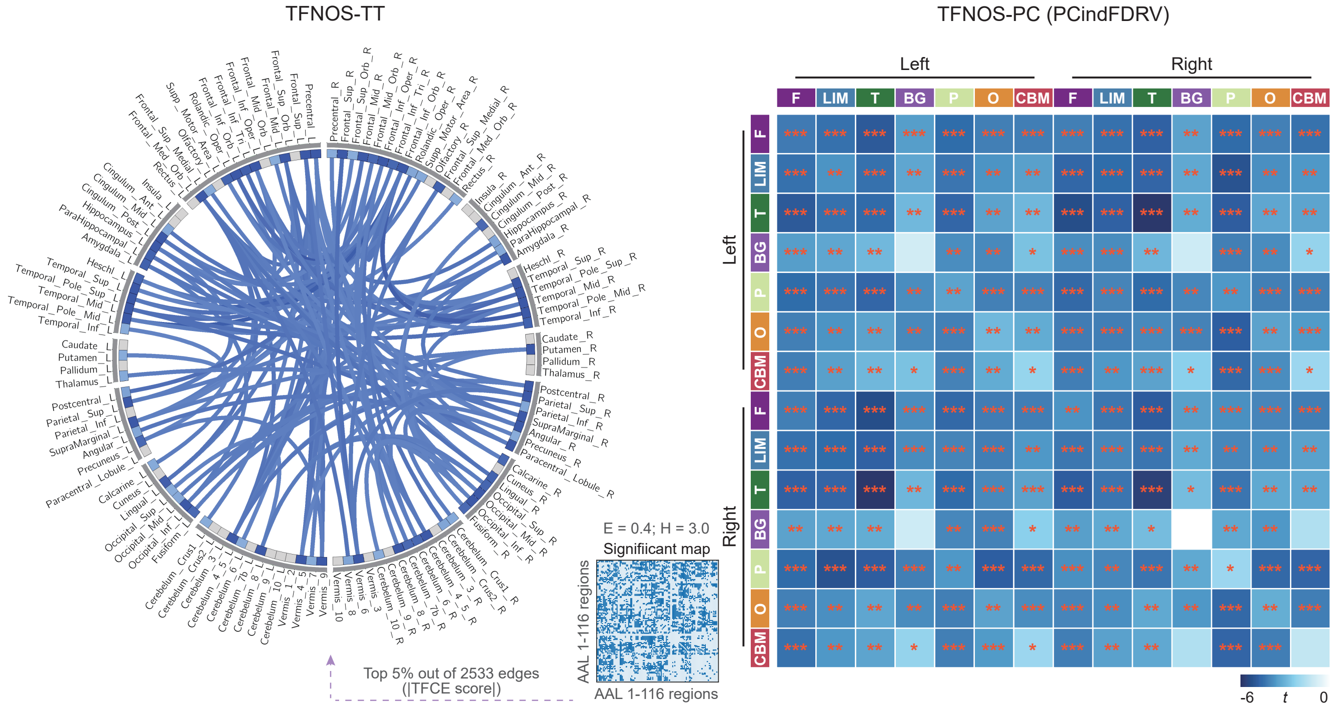


**Fig. S13.** Patients with SCI exhibit a widespread reduction in connectivity strength of Pearson-correlation-based undirected brain networks compared to HC. (Left) Using TFNOS-TT procedure with $E = 0.4$ and $H = 3.0$. For visualization purposes, the Circos plot only shows edges with $p < 0.05$ where the absolute value of the TFCE score is in the top 5% (out of the 2533 edges with $p < 0.05$). The blue lines indicate SCI < HC, with darker colors representing larger absolute TFCE scores. (Right) Using TFNOS-PC (PCindFDRV) procedure. $*p < 0.05$; $**p < 0.01$; $***p < 0.001$. F = frontal; LIM = limbic; T = temporal; BG = basal ganglia; P = parietal; O = occipital; CBM = cerebellum.

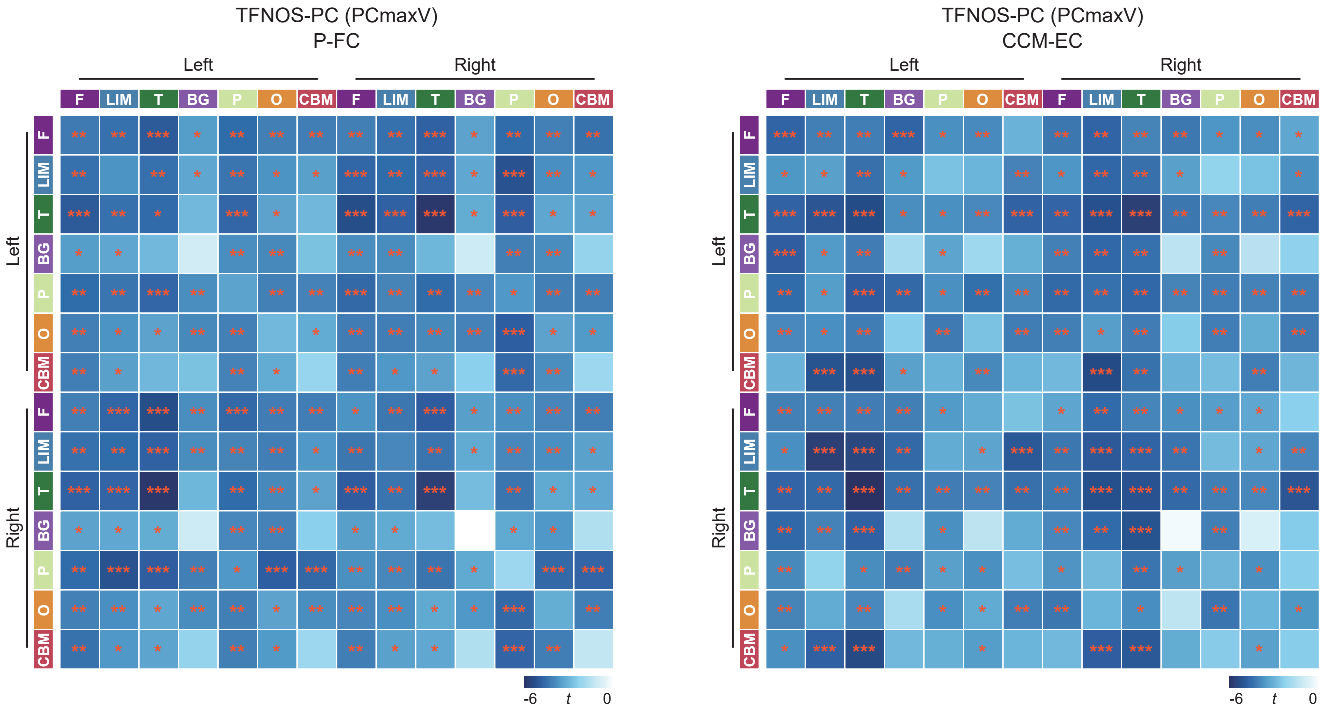


**Fig. S14.** TFNOS-PC (PCmaxV) procedure for examining differences between patients with SCI and HC. (Left) Using P-FC. (Right) Using CCM-EC. The PCmaxV is a strong-sense FWER-controlling procedure. $*p < 0.05$; $**p < 0.01$; $***p < 0.001$. F = frontal; LIM = limbic; T = temporal; BG = basal ganglia; P = parietal; O = occipital; CBM = cerebellum.
